# Fitness advantage of *Bacteroides thetaiotaomicron* capsular polysaccharide is dependent on the resident microbiota

**DOI:** 10.1101/2022.06.19.496708

**Authors:** Daniel Hoces, Giorgia Greter, Markus Arnoldini, Claudia Moresi, Sara Berent, Isabel Kolinko, Florence Bansept, Aurore Woller, Janine Häfliger, Eric Martens, Wolf-Dietrich Hardt, Claude Loverdo, Emma Slack

**Author notes:** Department for Evolutionary Theory, Max Planck Institute for Evolutionary Biology, Plön, Germany (Florence Bansept). Department of Molecular Cell Biology, Weizmann Institute of Science, Rehovot, Israel (Aurore Woller). Klinik für Gastroenterologie und Hepatologie, University Hospital Zurich, Zurich, Switzerland (Janine Häfliger).

## Abstract

Many microbiota-based therapeutics rely on our ability to introduce a microbe of choice into an already-colonized intestine. However, we remain largely blind to the quantitative effects of processes determining colonization success. In this study, we used genetically-barcoded *Bacteroides thetaiotaomicron* (*B.theta*) strains in combination with mathematical modeling to quantify population bottlenecks experienced by *B.theta* during gut colonization. Integrating population bottlenecks sizes with careful quantification of net growth rates *in vivo* and *in vitro* allows us to build models describing the events during intestinal colonization in the context of gnotobiotic and complex microbiotas. Using these models, we estimated the decrease in niche size for *B.theta* colonization with increasing microbiota complexity. In addition, our system can be applied to mechanistically dissect colonization defects of mutant strains. As a proof of concept, we demonstrated that the competitive disadvantage of a *B.theta* mutant lacking capsular polysaccharide is due to a combination of an increased lag-phase before growth initiation in the gut, combined with an increased clearance rate. Crucially, the requirement for the *B.theta* capsule depended strongly on microbiota composition, suggesting that the dominant role may be protection from bacterial or phage aggression rather than from host-induced bactericidal mechanisms.

## Introduction

From the moment that we first contact microbes at birth, we continuously encounter environmental and food-borne microbes. Whether such encounters are transient or will lead to long-term colonization, is influenced by complex ecological interactions between the invading species and the existing consortium, as well as the host’s dietary habits and the physiology of the intestine (David et al., 2014; Wotzka et al., 2019). A better understanding of the factors determining colonization efficiency is crucial in the development of microbiota engineering strategies (Donia, 2015; Pham et al., 2017; Sheth et al., 2016) and in the use of bacterial species as biosensors to probe microbiota function and stability (Goodman et al., 2009).

One established way of studying ecological processes within hosts is genetic barcode tagging of otherwise isogenic microbes, which has previously been used to study population dynamics of pathogens such as *Vibrio cholerae* or *Salmonella* Typhimurium within the infected host (Abel et al., 2015b; Vlazaki et al., 2019). Based on barcode tagging and mathematical modelling, it has been possible to infer parameters such as growth, clearance and migration rates (Dybowski et al., 2015; Grant et al., 2008; Kaiser et al., 2014, 2013), as well as the size of population bottlenecks imposed during colonization (Abel et al., 2015a; Li et al., 2013; Moor et al., 2017), antibiotic treatment (Vlazaki et al., 2020) or immunity (Coward et al., 2014; Hausmann et al., 2020; Lim et al., 2014; Maier et al., 2014; Moor et al., 2017). Furthermore, combining neutral genetic barcodes with targeted mutant strains, this experimental tool can be used to mechanistically analyze the fitness-effect of individual genes that regulate successful gut colonization or tissue invasion (Di Martino et al., 2019; Nguyen et al., 2020).

In order to study the dynamics of invasion of a novel microbiota member, we chose to use *Bacteroides thetaiotaomicron* (*B.theta*) as a model microbe. *B. theta* is a common commensal member of the human intestinal microbiota (Porter et al., 2018), and a well-established toolbox exists for engineering its genome (Porter et al., 2017; Whitaker et al., 2017). A common feature of *Bacteroides* species is the ability use phase variation to modulate the expression of 3-10 capsular polysaccharide (CPS) operons, leading to the production of distinct capsule structures (Porter and Martens, 2017). *B.theta* strains lacking a capsule have been shown to engraft poorly in an existing microbiota when competing with CPS-expressing strains (Martens et al., 2009; Porter et al., 2017). Critically for our studies, the deletion of all capsule gene clusters is not expected to negatively influence the growth rate of *B.theta* per se (Rogers et al., 2013) but is expected to affect survival on exposure to noxious stimuli, like bile acids, stomach acid, antimicrobial peptides, or phage (Porter et al., 2020, 2017), making this a good model to test our ability to quantify population dynamics *in vivo*. *B. theta* is also a relevant bacterium to understand population dynamics of colonization, as its ability to colonize to very high densities and the availability of precise tools for genetically engineering it (Lim et al., 2017; Mimee et al., 2015) make it a strong candidate for introducing novel functions into microbiomes.

To quantify the processes determining success of *B.theta* colonization in the presence of different resident microbiota, we generated genetically tagged *B.theta* strains with and without capsule. Using a combination of mathematical modeling and experimental quantification of population dynamics parameters, we estimated the probability of colonization success during single-strain colonization and competition. This revealed that acapsular *B.theta* have similar colonization success (i.e., encounter similar population bottlenecks) to wild-type strains when invading gnotobiotic microbiota communities, but not when invading a more complete microbiome. However, the acapsular strain still competes poorly with the wild-type strain in co-colonization even in gnotobiotic settings. The underlying mechanisms depend upon the increased clearance and extended lag-phase of the acapsular *B.theta in vivo*. Therefore, neutral tagging and mathematical modeling can be used to infer behavior of *B.theta* and protective functions of the polysaccharide capsule in a mouse model.

## Results

### Genetically-tagged *B.theta* strains to study within-host population dynamics

We first established tools that would allow us to quantify *B.theta* clonal loss *in vivo* and to test the plausibility of our theoretical models. Briefly, we used the previously described pNBU2 integration plasmid carrying an antibiotic resistance cassette, a phage promoter and an optimized ribosome binding site for fluorescent protein expression which allows visualization of clones (Wang et al., 2000; Whitaker et al., 2017). Six barcode tags, previously developed and validated for *Salmonella* (Wild-type Isogenic Tagged Strains, WITS (Grant et al., 2008; Maier et al., 2013) were inserted into an erythromycin-resistant (*ermG* cassette (Cheng et al., 2000) pNBU2 vector (Fig.1A). These six tags were inserted in both *B.theta* VPI-5482 *Δtdk*, which can phase vary the expression of eight different capsular polysaccharides (*B.theta* WT); and a *B.theta* strain that cannot produce capsule, generated by sequentially deleting all CPS gene clusters (*B.theta* acapsular) (Porter et al., 2017). A matched set of strains was produced in which an identical pNBU2-derived plasmid was integrated carrying the *tetQb* resistant cassette (Nikolich et al., 1992) in place of *ermG,* conferring instead tetracycline resistance (TetR). All strains also express a fluorescent protein, either GFP or mCherry (Fig.1A, Suppl.Table 1).

**Figure 1:**
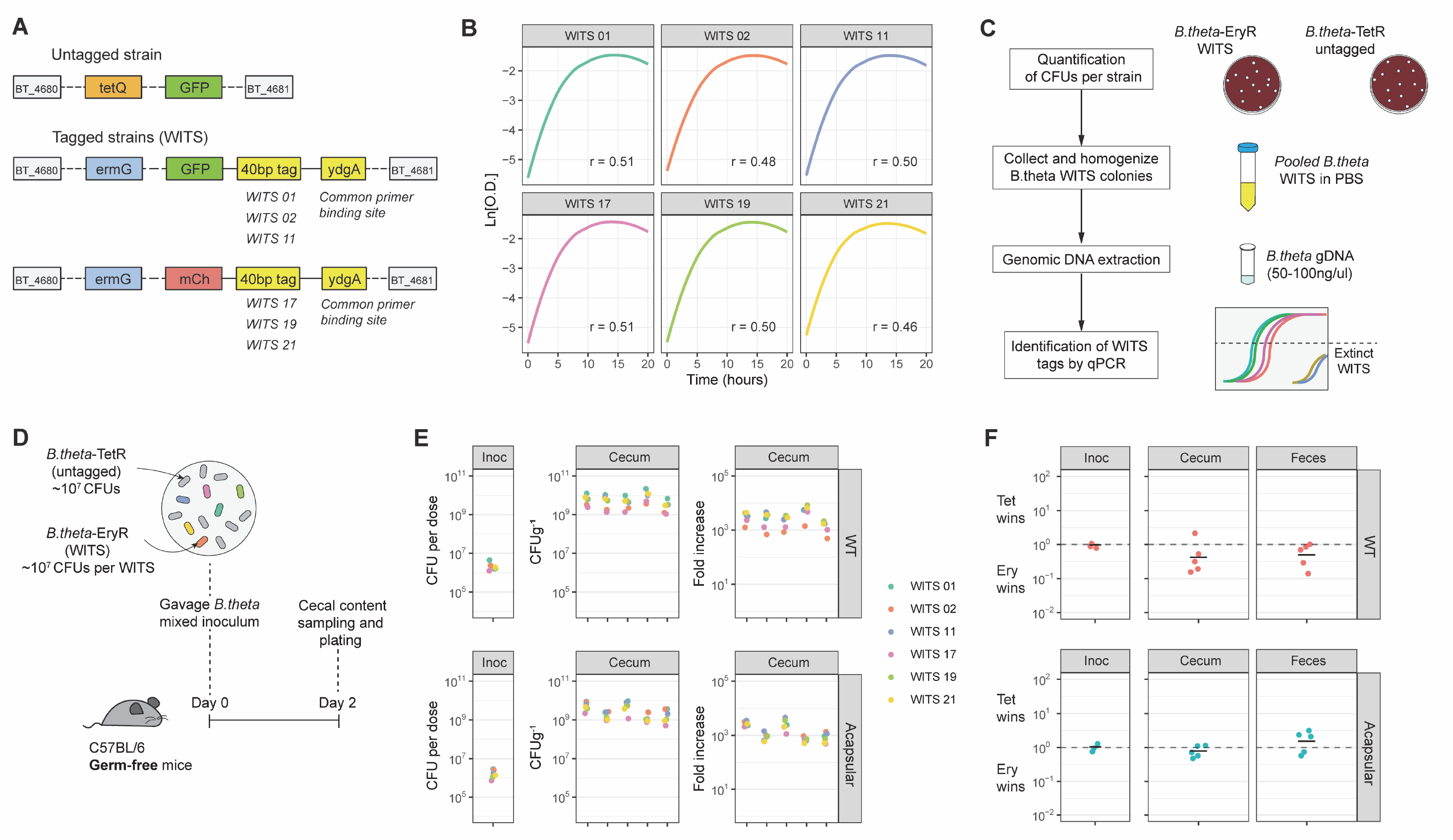
Tagged *B.theta* strains have similar fitness for growing *in vitro* and *in vivo*. **(A)** Schematic representation of insertions in *B.theta* genome. **(B)** Growth curves of *B.theta* WT in BHIS media (n=3 per strain) and growth rates (r, hours^-1^) per tag. **(C)** Workflow for sample preparation and data acquisition. **(D)** Experimental design of *in vivo* competitions. All strains (untagged *B.theta-*TetR and all six tagged *B.theta*-EryR WITS) were mixed in a 1:1 ratio. **(E)** Tags distribution during colonization among six *B.theta* tagged strains either WT or acapsular. Plots show distribution of tags in the inoculum, in cecal content of individual mice after 48h of colonization and fold increase of each tag per mouse compared to the inoculum. **(F)** Competitive index of tetracycline (Tet)- over erythromycin (Ery)-resistant strains *in vivo* after 48h of colonization in *B.theta* WT and acapsular in cecal content and in feces.

### Genetic barcode tags do not affect fitness of *B.theta* strains

A critical assumption of any analysis using genetically barcoded strains is that the chromosomal insertions, as well as the construction process, have not altered the fitness of the strains compared to the wild-type, both in culture and when colonizing a host. When grown individually in BHIS media, the *B.theta* strains carrying the different barcode tags had a similar growth rate (median doubling time ranged from 78-90 min) (Fig.1B and Suppl. Fig.1A). Also, they maintained their relative abundances, evaluated by qPCR (Fig.1C), when mixed and grown overnight (Suppl.Fig.1B and Suppl.Fig.1C).

To test whether the tags confer a fitness effect upon colonization of a host, we colonized germ-free mice with a uniform mixture of 10^7^ colony forming units (CFU) of each tagged strain (Fig.1D). At this tag abundance, stochastic loss of tags is highly unlikely. We compared the relative abundance of each tag in the inoculum to that in the cecal content and feces after 48h of colonization. These experiments revealed small, random deviations in the distribution, consistent with uniform fitness (Fig.1E). Finally, as erythromycin- and tetracycline-resistance were used to distinguish the tagged and untagged *B.theta* strains, we also tested whether the antibiotic resistant cassettes alter competitive fitness and found no significant difference between strains carrying the two resistance cassettes in culture (Fig.1F) or upon colonization (Suppl. Fig.1D).

### Determining inoculum size of tagged strains that yields maximal information upon ***B.theta* colonization**

We then applied the neutrally tagged *B. theta* strains to estimate their invasion probabilities by recovering *B. theta* cells after colonization and comparing the barcode distribution with that in the inoculum. As invasion probabilities depend on the interaction with the resident microbiota (Kurkjian et al., 2021), we probed *B.theta* colonization on three different communities: Low-complexity microbiota (LCM) (Stecher et al., 2010); Oligo Mouse Microbiota (OligoMM12) (Brugiroux et al., 2016) and specific-pathogen-free (SPF) complete microbiota (Suppl. Table 3). We evaluate recovering *B. theta* cells 48h after initial colonization, a timepoint shortly after the *B. theta* population in the cecum reaches carrying capacity (Suppl.Fig.2A).

Assuming that the change in relative abundance of tags before and after the colonization process is due to stochastic loss of *B. theta*, we formulated a mathematical model that describes this process and that allows us to infer important parameters, namely a per-cell colonization probability for *B. theta*. Mice are inoculated with *n_0_* copies of tagged *B.theta* cells, which undergo random killing during their transition through the stomach and small intestine. Surviving clones arriving in the cecum start growing and are detected 48h post-colonization (Fig.2A). A simple calculation was carried out to estimate the probability for the lineage of a bacterium clone spiked at *n_0_* copies in the inoculum to be found at the measurement time, which can be assumed as the combination of making it alive to the cecum and not being lost afterwards. As bacteria are not counted directly in the inoculum, but the inoculum is a sample of known volume of a solution of a known concentration, the incoming distribution of tagged bacteria is approximately Poisson, of mean *n_0_*. Then we assume each bacterium has an independent probability of colonization *β.* Therefore, the probability of losing an individual tag (which can be approximated to the fraction of tagged strains lost across different mice) is equal to *e*^-*βn*o^. Finally, the colonization probability (*β*) can be approximated to negative natural logarithm of the proportion of lost tags divided by the expected initial number of CFUs of each tagged strain in the inoculum. As there are experiments with different initial inoculum sizes (different *n_0_*), we obtain *β* by maximizing the likelihood of the experimental observations (See Methods, Mathematical modelling, section 1). Furthermore, a more complex calculation can be carried out using the variance of the tagged population rather than defined loss/retention (Suppl.Fig.2B and Suppl.Fig.2C; see Methods, Mathematical modelling, section 2), but the overall obtained numbers remain similar.

**Figure 2:**
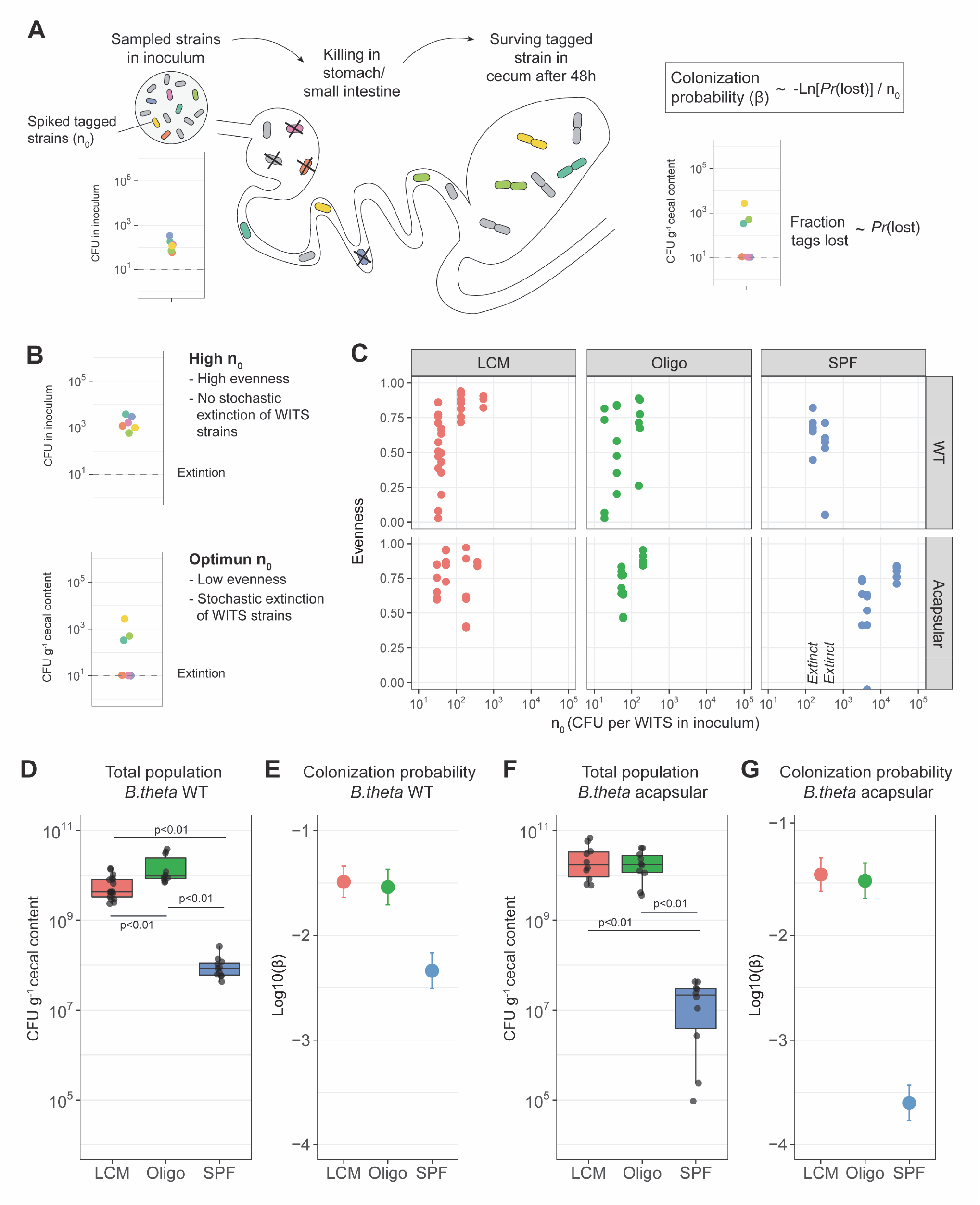
Colonization probability of *B.theta* strains in LCM, Oligo-MM12 and SPF mice. **(A)** Schematic representation of experimental estimation of colonization probability **(B)** Two different scenarios of *n_0_* and distribution of tagged strains: *n_0_* in which no extinction of tags is observed (“High *n_0_*”) and *n_0_* in which tags are randomly lost (“Optimal *n_0_*”). **(C)** Pielou’s evenness vs. *n_0_*. Pielou’s evenness was estimated with an maximum possible value of H_max_= ln(6) for all data points (six tagged strains). Each dot represents the evenness calculated per mouse. Values of *n_0_* for which all tags were lost, and therefore no evenness could be estimated, were marked with “Extinct” in the graph. **(D and F)** Total *B.theta* population in the cecum at 48h post-colonization for **(D)** WT and **(F)** acapsular strains. **(E and G)** Probability of colonization (β) in **(E)** WT and **(G)** acapsular in the cecum after 48h of colonization using the loss method. Estimation based on 6 tags times the total number of mice (LCM=17, OligoMM12=10, SPF=11) and on lost tags.

It is important to note that if all tags are lost, or if all tags are recovered, only upper or lower bounds for *β* can be determined, respectively, rather than numerical estimates. Thus, experiments where some, but not all tags are lost from the final population yield maximum information (Fig.2B). To find the inoculum sizes that lead to this outcome, we therefore titrated the tagged strains into an untagged *B. theta* population at different ratios. We used these mixtures to colonize mice carrying three different microbiota communities, and quantified tag recovery from the cecum 48h post-colonization. We used the Pielou evenness (Pielou, 1966) as a summary representation of the distributions of tagged population for both *B.theta* strains (WT and acapsular) in mice carrying three different microbiota communities (Fig.2C). After an analytical and numerical analysis, we found that the resulting estimates are most precise when about half the tags are lost (see Supplementary Methods). In the case of LCM and OligoMM12 gnotobiotic mice, an *n_0_* of between 10-100 was optimal for both wild-type and acapsular *B.theta*, whilst in SPF mice, an *n_0_*of around 500 was informative for wildtype, but 5000 CFU were needed of each acapsular tagged strain.

Finally, the observed change in the distribution of tags after the colonization process can have two explanations: stochastic loss of individual strains in this process, or mutation and selection in a subset of the strains. In order to test whether mutations, followed by selective sweeps, can explain observed changes, we recovered tagged *B. theta* strains from colonized mice, mixed them with the ancestral strains carrying different tags in equal ratios, and used this mix to inoculate a new group of mice. There was no consistent advantage of re-isolated strains over ancestral strains (Suppl.Fig.2D), indicating that mutation and selection do not play a role in the initial 48h after colonization.

### *B.theta* colonization probability in LCM, OligoMM12 and SPF mice

The resident microbiota composition in the mammalian gut is one of the main factors constraining the colonization of newly arriving species. This can happen through various mechanisms such as competition for nutrients (Baumler and Sperandio, 2016; Maier et al., 2013), modification of the intestinal environment (Cremer et al., 2017) or via direct suppression of the invaders by phages (Almeida et al., 2019; Barr et al., 2013) or Type VI secretion systems (Chatzidaki-Livanis et al., 2016; Wexler et al., 2016). Therefore, we used the previously described mathematical model to estimate the suppressive potential of different microbiotas, gnotobiotic (LCM, OligoMM12) and SPF, on *B. theta* invasion.

*B.theta* load in the cecal content after 48h of post-inoculation were similar in LCM and OligoMM12 mice, but significantly lower in SPF mice (Fig.2D). We then used the measured tag loss to determine colonization probabilities as described above, for the different *B. theta* strains in the different microbiota backgrounds. The colonization probability of *B.theta* WT was lower in SPF mice (Log10(*β,* colonization probability) ± 2 standard deviations = −2.35 ± 0.14) compared to the two gnotobiotic microbiotas (LCM, −1.50 ± 0.10; and Oligo, −1.54 ± 0.13) (Fig.2E). This additionally reveals that there is a discrepancy between the relative size of the final population (100-fold lower in SPF mice) and the relative probability to recover a specific clone from the inoculum at 48h (only 10-fold lower in SPF mice), indicating that size of the open niche does not linearly translate into colonization probability. For these two observations to agree, the founding population of *B.theta* has to be smaller in SPF mice, and the net growth during the first 48h post-inoculation has to be lower than in the gnotobiotic mice (Suppl.Fig.2A).

As capsular polysaccharides (CPS) are thought to play a crucial role in phage evasion/infection (Porter et al., 2020), immune evasion (Fanning et al., 2012; Hsieh et al., 2020; Porter et al., 2017) and protection from other environmental stressors, we expected to see a tighter bottleneck (decreased colonization probability) for the acapsular *B.theta* strain in all microbiota backgrounds. Surprisingly, in gnotobiotic mice, the total population size of acapsular *B.theta* and the probability to colonize (Log10*β*: LCM, −1.43 ± 0.13; and OligoMM12, −1.49 ± 0.14) were not different to the WT strain (Fig.2F and Fig.2G). Thus, there is no measurable fitness benefit of CPS in gut colonization up to 48h post-inoculation in these settings. However, we observed a different scenario when we inoculated acapsular *B.theta* into mice carrying a SPF microbiota. Both the total population size of acapsular *B.theta* (Fig.2F) and the colonization probability (Log10*β*: SPF, −3.65 ± 0.13; Fig.2G) were 10-fold lower compared to the wildtype strain, indicating a fitness benefit of CPS in the context of a more diverse microbiota and a smaller niche. Given the previously observed absence of a growth defect for acapsular *B.theta*, we hypothesized that increased clearance of acapsular *B. theta* cells causes the difference, potentially attributable to competition/aggression from other microbiota species or from more active host immunity in SPF mice.

### Competitive colonization with acapsular and WT *B.theta* reveals a role of CPS in gnotobiotic mice

The absence of an increased bottleneck during colonization of gnotobiotic mice with acapsular *B.theta* is in conflict with previous studies showing a fitness defect of this strain (Coyne et al., 2008; Porter et al., 2017). We therefore carried out competitive colonization with *B.theta* WT and acapsular *B.theta*. Starting at a 1:1 ratio we inoculated mice with erythromycin-resistant WT and tetracycline-resistant acapsular *B.theta* and quantified the cecal bacterial load 48h post-inoculation (Suppl.Fig.3A). This reveals a gradient of outcompetition of the acapsular strain ranging from a competitive index (abundance of WT over acapsular *B.theta* at the end of the experiment) of approx. 20 in germ-free mice, 100 in gnotobiotic mice and 10^4^ in SPF mice (Fig.3A). This demonstrates a competitive fitness benefit of CPS, that increases in proportion to microbiota complexity.

**Figure 3.**
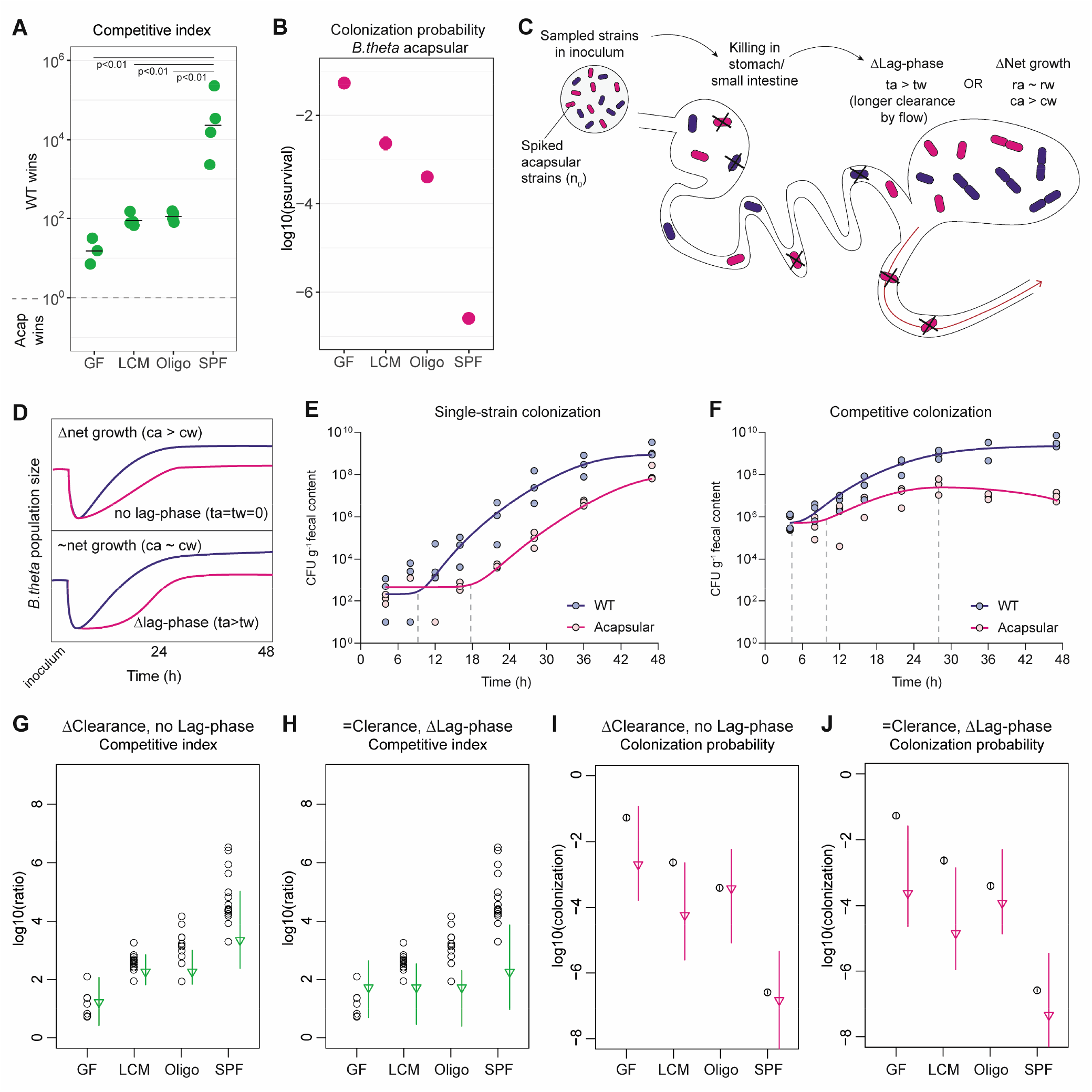
Competition colonization with acapsular and WT strains. **(A)** Competition index (ratio between WT over acapsular in the cecum) after 48h of colonization starting at a 1:1 ratio. **(B)** Probability of colonization by the acapsular strain during competition with the WT strain. Tagged acapsular strains were spiked into a WT-strain inoculum (GF=7, LCM=13, OligoMM12=12, SPF=16). **(C)** Schematic representation of two scenarios of competitive advantage of the WT over the acapsular *B.theta* having a similar initial probability of colonization of the cecum: difference in lag-phase (mean time to growth commencement in acapsular (ta) and WT (tw)) and difference in net growth rate (growth rate in acapsular (ra) and WT (rw); clearance rate in acapsular (ca) and WT (cw)). Clearance can be due to both flow/loss in the fecal stream and death. **(D)** Growth curves representing the two scenarios in (C). **(E)** Fecal bacterial load of *B.theta* WT or acapsular during single-strain colonization in OligoMM12 mice. Overlay graph of individual experiments with starting inoculum 10^2^-10^3^ CFU each. **(F)** Fecal bacterial load of *B.theta* WT or acapsular during competitive colonization in OligoMM12 mice. Overlay graph with starting inoculum 10^6^ CFU each (ratio 1:1). **(G-J)** Estimation of the competitive index and colonization probability of the acapsular strain (see Supplementary Methods): **(G and I)** assuming no lag-phase and estimated difference in clearance rate between the WT and acapsular strains and **(H and J)** assuming a mean 6h difference in lag-phase and identical clearance rate between the WT and acapsular strains (circles: experimental data; triangles: mean parameter estimation and error).

To confirm these findings, we performed a competition experiment in the same microbiota backgrounds, but this time, using tetracycline-resistant *B.theta* WT and barcoded erythromycin-resistant acapsular *B.theta* strains. WT strain density (Suppl.Fig.3B) and the average increase in the WT relative to the acapsular strains (after adjusting for the initial ratio in the inoculum, Suppl.Fig.3C) were similar between the two experiments. Interestingly, when calculating the colonization probability *β* based on tag loss as described before, this is lower for the acapsular *B. theta* strain when co-colonizing with the WT strain than when colonizing alone (Fig.3B). This indicates that the competition with wild-type results in both a lower total population size and increased clonal extinction in the acapsular strains.

### Longer lag-phase and higher clearance rate explains fitness defect of *a*capsular *B.theta* in competitive colonization

In order to understand how acapsular *B.theta* can have an indistinguishable colonization probability when inoculated alone, but a major fitness defect in competition with wildtype *B.theta,* we extended our simple, one-step colonization model to include a lag-phase after arrival in the cecum (Fig.3C). Based on this model, two different but non-exclusive scenarios can explain the competitive disadvantage of the acapsular strain during competition. In the first scenario, there is a difference in the net growth rate, therefore explaining the ratio between acapsular and WT strains at the end of the experiment (Fig.3D, top). In the second scenario, both strains have similar net growth rates but a different lag-time, which delays initial growth of the acapsular such that the wild-type will occupy a larger fraction of the available niche during competition (Fig.3D, bottom).

To evaluate these two different scenarios, we carried out a time-course analysis of the fecal bacterial load of *B.theta* during colonization in OligoMM12 mice for the acapsular and WT strains alone and in competition. This allows us to extract the *in vivo* net growth rates, i.e., growth rate minus clearance rate, as well as lag-phases between strains. Interestingly, the acapsular strain does exhibit an initial lag-phase of approximately 4-8h when compared to the WT, even in single colonization (Fig.3E and Fig.3F). Based on the CFU data alone, this delay in colonization could either be attributed to a tighter population bottleneck during small intestinal transit, or to a longer lag-phase of clones of the acapsular strain after arrival in the large intestine, or potentially to both. However, since our neutral tagging data indicates an identical bottleneck of wild-type and acapsular *B.theta* strains when colonized individually in OligoMM12 (Fig.2E and Fig.2G), we can exclude increased killing of the acapsular strain prior to seeding of the cecum. Once in the cecum, the net growth rate during the exponential growth phase is lower for the acapsular strain (WT: 0.53±0.05 and acapsular: 0.40±0.02; estimated from Fig.3E). As the growth rates of these strains in culture is identical, it is reasonable to assume that the slightly lower net growth rate is due to an increased clearance rate for the acapsular *B.theta* in the gut lumen. This is further supported by the observed decrease in the total population of the acapsular strain after the WT has reached carrying capacity (Fig.3F).

We then use these experimentally-measured parameters to theoretically estimate the competitive index and the colonization probability of the acapsular strain during competition. We modelled three different scenarios assuming either a difference in clearance rate or lag-phase, or a combination of both (see Supplementary Methods for the description of all parameters used). Comparison of the modeled values with the observed data will then indicate whether the model can capture the dominant part of the phenomenon. We found that the competitive index (abundance of WT over acapsular strain) can be mostly explained by a model in which only clearance rate differs between strains (Fig.3G). It was not possible to fit a model in which only the lag-phase differed for the observed competitive index (Fig.3H). In contrast, models based either on an increased clearance rate (Fig.3I) or on a prolonged lag-phase (Fig.3J), could explain the observed colonization probability of the acapsular during competition. Combining both parameters showed a similar tendency to the initial model without a lag-phase (Suppl.Fig.3D and Suppl.Fig.3E). Therefore, while we clearly observe a delayed start of growth in the acapsular strain in vivo, the increased clearance rate clearly has a stronger effect on the overall abundance of the acapsular strain during competitive colonization.

### Acute challenges modify *B.theta* population dynamics in vivo

Finally, as a proof of concept for neutral tagging in the study of established microbiota, we used our system to probe clonal extinction when an established *B.theta* population in the gut is challenged by two major environmental perturbations: 1) Shifting from standard chow to a high-fat no-fiber diet (HFD) (David et al., 2014; Wotzka et al., 2019)) and 2) infectious inflammation driven by *Salmonella* Typhimurium (Stm) (Maier et al., 2014; Wotzka et al., 2017).

In order to have a highly simplified system to study host-*B.theta* interactions, we monocolonized germ-free C57BL/6J mice with WT *B.theta* spiked with tagged strains in such a way to maintain all tags with a roughly uniform distribution, i.e., minimum loss, in the gut lumen prior to challenge (Fig.4A). After 4 days of colonization, we exposed mice to either HFD or oral infection with 10^6^-10^7^ CFUs of attenuated Stm SL1344^ΔSPI-2^. This attenuated strain was used to induce moderate intestinal inflammation in the gnotobiotic mice but to avoid the rapid lethal infections associated with wildtype *Salmonella* in germ-free animals (Hapfelmeier et al., 2005; Stecher et al., 2005). *B.theta* populations were monitored in feces before the challenge was administered and three days after the challenge in feces and cecum.

**Figure 4.**
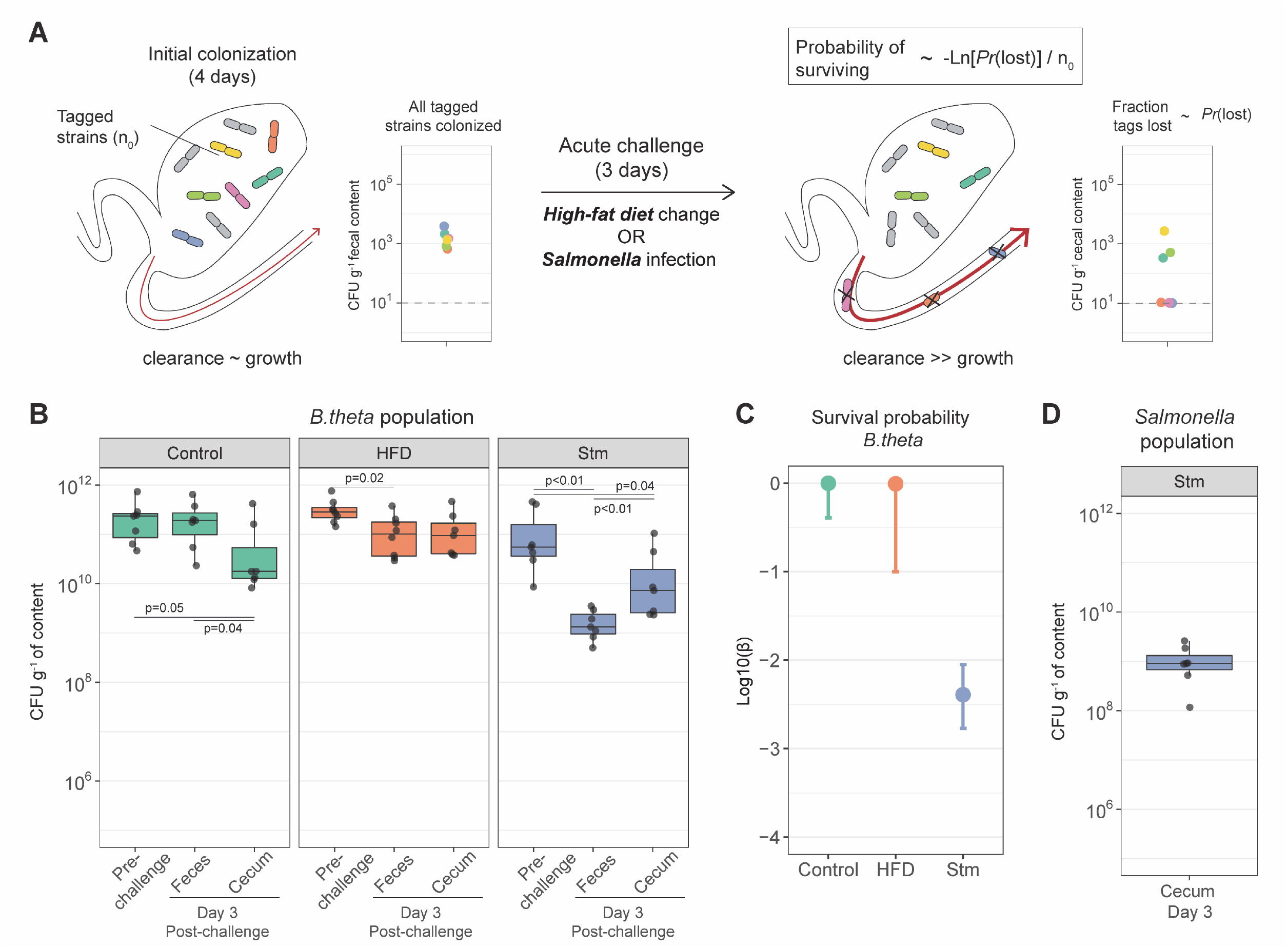
Acute challenges imposed a population bottleneck in the resident *B.theta* population. **(A)** Schematic representation of experimental estimation of colonization survival probability after acute challenges. **(B)** Population of *B.theta* in monocolonized ex-GF mice kept under standard chow (control) and during acute challenge with high-fat diet (HFD) or attenuated *Salmonella* infection. **(C)** Probability of surviving in the cecum after 3 days of the acute challenge. Estimation based on 6 tags timer the total number of mice (Control=7, HFD=8, Stm=7). **(D)** Population of attenuated Salmonella in cecal content.

Despite published reports that *B.theta* is sensitive to bile acids (Wotzka et al., 2019), HFD did not significantly change the cecum population size (Fig.4B) or increased loss of tagged *B.theta* (Fig.4C) in the diet-shift experiments. We can still set a lower-bound for the probability of an established *B.theta* clone to survive the acute dietary challenge by estimating the value in case that one tagged strain was lost during the challenge. In this case, HFD imposed a similar survival probability (Log10(*β*) = −1 to 0) to the one observed in the control group (Log10(*β*) = - 0.39 to 0). Estimating that the initial *B.theta* population is in the order of 10^10^ CFUg^-1^ in the cecum, the bottleneck effect imposed by HFD would reduce the population minimally to a range between 10^9^ to 10^10^ clones, which are densities normally observed in the cecum (Fig.4B).

In contrast, after three days of infection, *Salmonella* induced-inflammation reduced the total *B.theta* population approx. 100-fold in feces at day 3 post-infection (Fig.4B). In addition, we estimated the probability of an established clone to survive this acute inflammation challenge is approximately 1 out of 500 in cecum (Fig.4C) while competing with the *Salmonella* population (Fig.4D). This means that the initial estimated population of 10^10^ CFUg^-1^ in the cecum is reduced to a population of between 10^7^ to 10^8^ clones by the bottleneck effect induced by inflammation and then bouncing back to the observed levels between 10^9^ to 10^10^ CFUg^-1^ (Fig.4B). Therefore, the bottleneck effect on the population induced by acute inflammation not only reduces the total population size but also diminishes the diversity of clones in the population.

## Discussion

Understanding the different mechanistic factors determining how microbes can invade a resident gut microbiota is of considerable importance for combating mucosal infections (Kreuzer and Hardt, 2020; Stecher, 2021), but also for rationally introducing new functions into existing communities (Cubillos-Ruiz et al., 2021). These factors can include, among others, nutrient/energy availability, environmental factors such as pH and flow/dilution rate (Arnoldini et al., 2018), and the presence of directly toxic or aggressive activities derived from the host (Cullen et al., 2015) or from other microbiota species (García-Bayona and Comstock, 2018), all of which can affect different microbes in different ways. Here, we condense all these mechanisms into three processes: factors affecting immigration rate (i.e., arrival into growth-permissive sites in the gut), factors affecting growth, and factors affecting clearance/death (Hoces et al 2020). Combining well-designed experiments using genetically barcoded strains with mathematical modeling, we were able to empirically (e.g., net growth rates through plating) or deductively (e.g., probability of colonization) estimate the relative importance of these three processes for colonization of *B. theta* under different conditions. In addition, we were able to study mechanistically how fitness-relevant genetic changes in *B. theta* (production of capsular polysaccharide) affect colonization success.

When analyzing the effect of capsular polysaccharide expression in the process of colonizing mice with different resident microbiotas, we found that the previously shown fitness disadvantage of acapsular strains (Coyne et al., 2008; Porter et al., 2017) depends on the microbiota context, rather than on host effects. Both, the wild type and the acapsular strains colonize mice with a relatively simple microbiota (LCM, Oligo) similarly well. However, in mice with more complex microbiota (SPF), the acapsular strain engrafted significantly less well than the wild type. While a possible explanation includes more robust intestinal immunity in fully colonized SPF mice, the simplest explanation is that expression of CPS is primarily important for interaction with or protection against other microbes. Possible microbe-inflicted processes against which CPS can protect include microbe-on-microbe killing (Chatzidaki-Livanis et al., 2016; Wexler et al., 2016) and susceptibility to phages (De Sordi et al., 2019; Porter et al., 2020). Co-colonization experiments with mixed *B. theta* inoculums consisting of wild type and acapsular strains revealed the specific disadvantages of the acapsular strain: a longer lag-phase before it starts growing in the gut and a higher clearance rate.

The effect of two important challenges with known effects on resident commensals, high-fat feeding (Wotzka et al, 2019) and inflammation (Maier et al, 2014), differentially affect *B. theta* and seem to depend on the ecological context in different ways. Feeding mice which are stably monocolonized with *B. theta* a fiber-less high-fat diet imposes a surprisingly mild bottleneck on the *B. theta* population 3 days after treatment, even though this intervention has been shown to increase bile acid concentrations to levels that inhibit *B. theta* growth (Wotzka et al, 2019). As it has been shown that *B.theta* rapidly evolves to adapt to dietary challenges in the context of a resident microbiota (Dapa et al., 2022), the observed mild population bottleneck imposed by high-fat diet feeding might only manifest if *B. theta* competes against other gut microbiota members, potentially ones that are more resistant to bile salts (e.g., *E. coli*, see Wotzka et al, 2019). When infecting mice that are stably monocolonized with *B. theta* with Salmonella, we observed a larger decrease in *B. theta* clonal survival probability, which is consistent with the sensitivity of commensal species to gut inflammation (cite Stecher et al, 2007). Interestingly, inflammation also causes a population bottleneck in the infecting *Salmonella* population (Maier et al, 2014), but it is less pronounced than the one we observe for *B. theta*, and the total population size of *Salmonella* rapidly recovers after this bottleneck. Therefore, the rapid killing/clearance of gut luminal *B. theta* seems to be representative of microbiota suppression that underlies the loss of colonization resistance observed in Salmonella-induced colitis.

Of course, our mathematical models are based on certain assumptions which are useful to simplify calculations but always risk introducing bias. Most notably, we have made estimates for a single population of bacteria that has a constant growth and clearance rate. Necessarily the reality is more complex than this – the nutrient profile and motility of the intestine will vary with circadian rhythm. Also previous work has demonstrated that particular CPS-expressing clones may have an advantage colonizing the mucus (Donaldson et al., 2018), although gnotobiotic studies suggest that at the population level *B.theta* grows at a similar rate in the mucus layer and lumen (Li et al., 2015). Nevertheless, recognizing these limitations, our estimates of colonization probability, growth and clearance rates still give a good overview of the harsh processes with strong effects on the *total* intestinal *B.theta* population. Future works should include individual-based models that can evaluate the impact of bacterial clones with different distributions of growth/clearance rates, as well as working with experimental models (microfluidics, for example) that would allow experimental investigation of the impact of single-cell level variation on total population behavior.

By combining mathematical modelling with direct quantification of bacterial population dynamics, we can gain insight into the major phenomena influencing colonization efficiency. Not only do these results help us understand the different steps of *B. theta* colonization, but they also serve as a proof of concept for studying other complex, multi-step biological processes using the set of experimental and data-analysis tools we are describing here. This raises the possibility to optimize colonization conditions in order to promote the efficient engraftment of beneficial species into target microbiota, or to better understand the processes of invasion of pathogens and the functional basis of colonization resistance.

## Acknowledgements

We express our gratitude to Sven Nowok and Dominik Bacovcin for their great support in maintaining the gnotobiotic mouse facility and to the whole team in EPIC and RCHCI. Also, we thank Dr. Annika Hausmann and Verena Lentsch for the discussions about this project. We acknowledge Prof. Bärbel Stecher for provision of the OligoMM12 bacterial strains used to generate the mouse colony used in these experiments.

## Funding

This work was funded by NCCR Microbiomes, a research consortium financed by the Swiss National Science Foundation (E.S., W-D.H); Swiss National Science Foundation (40B2-0_180953, 310030_185128) (E.S.), European Research Council Consolidator Grant (NUMBER 865730-SNUGly) (E.S.), Gebert Rüf Microbials (GR073_17) (E.S., C.L.); Botnar Research Centre for Child Health Multi-Invesitigator Project 2020 (BRCCH_MIP: Microbiota Engineering for Child Health) (E.S.), Agence Nationale de la Recherche (ANR-21-CE45-0015, ANR-20-CE30-0001) (C.L.) and MITI CNRS AAP adaptation du vivant à son environnement (C.L.). The funders had no role in study design, data collection and analysis, decision to publish, or preparation of the manuscript.

## Author Contributions

Conceptualization, D.H., M.A., C.L. and E.S.; Methodology, D.H., G.G., M.A., I.K., F.B., A.W., E.M., C.L. and E.S.; Formal Analysis, D.H., M.A., C.L., and E.S.; Investigation, D.H., G.G., C.M., S.B., C.L.; Resources, I.K., J.H., E.M., W-D.H., C.L. and E.S.; Writing – Original draft, D.H. and E.S.; Writing – Review and Editing D.H., G.G., M.A., C.M., S.B., I.K., F.B., A.W., J.H., E.M., W-D.H., C.L. and E.S.; Visualization, D.H., M.A., C.L. and E.S.; Supervision, C.L. and E.S.; Funding acquisition, C.L. and E.S.

## Declaration of Interest

The authors declare no competing interests.

## Methods

### Bacterial strains and cultures

B. theta strains were grown anaerobically (5%H_2_, 10%CO_2_, rest N_2_) at 37°C, overnight in brain heart infusion (BHI) supplemented media (BHIS: 37g/L BHI (Sigma); 1g/L-cysteine (Sigma); 1mg/L Hemin (Sigma)). For enrichment cultures in plates, we used BHI-blood agar plates (37g/L BHI (Sigma); 1g/L-cysteine (Sigma); 10% v/v defribinated sheep blood (Sigma)). Antibiotics were added to liquid cultures or plates as required for strain selection: erythromycin 25µg/L or tetracycline 2µg/L. In the case of BHI-blood agar plates used for cloning or gut content enrichment, we additionally added gentamycin 200µg/L to all plates. Plates were incubated for 48-72h at 37°C in anaerobic conditions. For a complete list of the bacterial strains used in this study, see Suppl. Table 1.

### Isogenic tag construction and integration

Genetic tags, fluorescent proteins and antibiotic resistance were introduced by using the mobilizable *Bacteroides* element NBU2, which integrates into the *Bacteroides* genomes at a conserved location (Wang et al., 2000). Gene fragments containing a unique 40bp sequence (biding site for forward primer) and a 609bp sequence with the ydgA pseudogene (common binding site for reverse primer) were synthesized (gBlocks, Integrated DNA Technologies) and cloned by Gibson Assembly Master Mix (NEB) into an NBU2 plasmid carrying the erythromycin resistant cassette ermG (tagged *B.theta*-EryR strains). A similar NBU2 plasmid carrying the tetracycline resistant cassette tetQb was used to construct the untagged strains (*B.theta*-TetR). For both, we used 10µL of desalted assembly reaction products to transform competent *E.coli* S17-1 cells (mid-log cells, washed three times in deionized ice-cold water) by electroporation (V=1.8kV; MicroPulser, BioRad). After 1h recovery at 37°C in 1mL of LB, cells were plated on LB plates with chloramphenicol (12µg/mL) and grown overnight. Plasmid-carrying *E.coli* S17-1 and *B.theta* strains were cultured overnight in 5mL of liquid media. *E.coli* S17-1 and *B.theta* were washed with PBS, pooled in 1mL of PBS, plated BHI-blood agar plates without antibiotics, and grown aerobically at 37°C for at least 16 hours. The lawn of *E.coli* S17-1 and *B.theta* was collected in 5mL of PBS, homogenized by vortex and 100µL were plated in BHI-blood agar plates supplemented with erythromycin 25µg/L and gentamycin 200µg/L. After 48 hours, single colonies were streaked in fresh BHI-blood agar plates with antibiotics, to avoid potential contamination with WT strains. Successful insertion in the BTt70 or BTt71 sites was evaluated by PCR.

### *In vitro* growth curves and competition

Individual *B.theta* strains were grown overnight on BHIS. Stationary-phase cultures were washed with PBS, and O.D. was quantified and adjusted to 0.05 in 200µL of fresh BHIS in a round 96-well tissue culture plates. Plates were transferred into the anaerobic tent and growth was quantified at 37°C with shaking, using a plate reader (Infinite PRO 200, Tecan).

For competition experiments, stationary-phase cultures *B.theta* WT or acapsular strains were washed with PBS, O.D. was quantified and adjusted to approximately 5×106 CFU/mL per strain (one *B.theta*-TetR and six *B.theta*-EryR tagged strains) in 10mL of fresh BHIS. An aliquot of this mix was serially diluted and plated in BHI-blood agar plates with the respective antibiotics for CFU quantification. Cultures were kept overnight, with shaking (800rpm) at 37°C in the anaerobic tent. Afterwards, an aliquot was plated as described before. For assessing the competition between *B.theta* tagged strains, we isolated DNA from one of the dilutions used for quantification and assessed the relative distribution of the tags by qPCR (see Quantitative PCR section). For assessing the competition among strains with different antibiotic resistances, we calculated the competition index by dividing the CFU/mL of the untagged *B.theta*-TetR strain (tetracycline-resistant) by the adjusted number of *B.theta-*EryR tagged strains (erythromycin-resistant; CFU/mL divided by six, as all the tagged strains were present in the culture).

### Mice

All animal experiments were performed with approval from the Zürich Cantonal Authority. In all experiments, we used mice with C57BL/6J genetic background, between 12-15 weeks old and of variable gender. C57BL/6J germ-free and gnotobiotic mouse lines (Low-complexity microbiota (LCM) (Stecher et al., 2010); Oligo Mouse Microbiota (OligoMM12) (Brugiroux et al., 2016)) were raised in surgical isolators under high hygiene standards at the ETH Phenomic Center. C57BL/6J SPF mouse line was also raised in the same facility. Mice were transferred to our experimental facility in sterile, tight closed cages and house into the IsoCage P-Bioexclusion System (Tecniplast) for 24h-48h before the experiment to adapt to new housing conditions. In all experiments, standard chow and water was supplemented under strict sterile conditions to avoid potential contaminations.

### *In vivo* growth curves and competition

B.theta WT WITS 01 strain was grown overnight on BHIS. Stationary-phase cultures were washed with PBS, and an inoculum of ∼5×10^7^ CFUs/100µL dose was prepared. C57BL/6J mice carrying the described microbiota composition (LCM, OligoMM12, SPF, 4 mice per group) were gavaged with the inoculum and fecal pellets were collected approximately every 4h for the first 20h, followed by two extra time points at 29h and 48h post-inoculation. Fecal pellets were weighted and homogenized in 500µLof PBS with steel ball by mixing (25Hz, 2.5min) in a TissueLyser (Qiagen). Serial dilutions were plated for quantification on BHI-blood agar plates supplemented with gentamycin 200µg/L and erythromycin 25µg/L.

For competition experiments, *B.theta* strains were grown overnight in 8mL of BHIS with corresponding antibiotics (erythromycin 25µg/L or tetracycline 2µg/L). Each culture was spun down at 3000g for 20min and resuspended in 10mL of PBS and individual O.D. was measured (cell number estimation 1 O.D.=∼4×10^8^ cells/mL). Each strain was adjusted to approximately 5×10^6^ CFU/100µL dose per strain in the inoculum mix (one *B.theta*-TetR and six *B.theta*-EryR tagged strains). Germ-free mice were gavaged with 100µL of the inoculum. After 48h, fecal and cecal contents were collected. Fecal content was homogenized and plated for quantification as described before. Cecal content was resuspended in 1mL of PBS and homogenized with steel ball by mixing with the same protocol (25Hz, 2.5min). Serial dilutions were prepared and plated in BHI-blood agar plates supplemented with gentamycin plus either erythromycin or tetracycline for CFU quantification of each strain. Similar to the in vitro competition experiment, we isolated DNA from one of the dilutions used for quantification to assess the competition between *B.theta* tagged strains. Relative distribution of the tags was obtained by qPCR. For calculating the competition index among strains with different antibiotic resistances, we divided the bacteria density of the untagged *B.theta*-TetR strain by a sixth of the bacteria density of the total *B.theta*-EryR tagged strains (as all six tagged strains were present in the culture).

### Colonization experiments

Stationary-phase *B.theta* strains were grown overnight as described before. Each culture was washed once with PBS and adjusted in the inoculum based on its O.D. For the untagged strain (*B.theta-*TetR), ∼5×10^7^ CFUs/100µL dose were transferred to the inoculum. For the *B.theta*-EryR tagged strains, we prepared an initial mix of all tagged strains in 50mL of PBS to a concentration of 10^5^ CFU/mL of each strain. After mixing by vortex for 1min, the required amount of *B.theta-*EryR tagged strains was spiked into the inoculum (between 30 - 5×10^4^ CFUs depending on the experiment). Gnotobiotic (LCM, Oligo) and SPF C57BL/6J mice were gavaged with the 100µL inoculum. Inoculum was serially diluted and plated for quantification of CFUs in BHI-blood agar plates supplemented with gentamycin 200µg/L plus either erythromycin 25µg/L or tetracycline 2µg/L. In addition, 2-3 doses (100µL) were directly plated in BHI-blood agar plates with gentamycin plus erythromycin to address initial distribution of *B.theta*-EryR tagged strains in the inoculum by quantitative PCR (qPCR). Two days after colonization, mice were euthanized and cecal content was collected in 2mL Eppendorf tubes and weighted. Cecal content was homogenized as described before. Serial dilutions were prepared and plated in BHI-blood agar plates supplemented gentamycin plus either erythromycin or tetracycline for CFU quantification. In addition, 100µL of homogenized content was plated directly in BHI-blood agar plates with gentamycin plus erythromycin for further assessment of the distribution of *B.theta*-EryR tagged strains by qPCR.

### In vivo competition of post-colonization versus original strains

To discard potential increased colonization fitness in the tagged strains that were present in the cecum content after 48h, we isolated single *B.theta* WT tagged strains that were present in the cecal content of SPF during a colonization experiment. Single colonies were expanded in liquid media, and the presence of a single strain was confirmed by qPCR analysis of the tagged tag. We randomly selected three of the *B.theta* WT tagged strains isolated from the cecal content. We prepared an inoculum as described before for the in vivo competition experiments with approximately 5×10^6^ CFU/100µL dose per strain in the inoculum mix. We complemented the inoculum with the remaining three *B.theta* WT tagged strains coming from the original stock. SPF mice were inoculated by gavage and cecal content was collected 48h later. Cecal content was processed as described before for CFU quantification and relative tag distribution by qPCR.

### Diet modification and Infection challenge experiments

In accordance with what we described before, *B.theta* WT strains were grown overnight in BHIS with corresponding antibiotics. As the untagged strain *B.theta*-TetR was used in higher concentrations, we prepared between 50-100mL of liquid culture depending on the number of mice to colonize. Inoculum was prepared as previously described with a concentration of 10^8^- 10^9^ CFUs/100µL dose of untagged *B.theta*-TetR, spiked with approximately 30 CFU of each *B.theta*-EryR tagged strains. Germ-free mice were gavaged with 100µL of the inoculum. Mice were maintained on standard chow diet (Kliba Nafag, 3537; autoclaved; per weight: 4.5% fat, 18.5% protein, ∼50% carbohydrates, 4.5% fiber) for 4 days. Afterwards, mice were housed on fresh IsoCages and challenges were applied as follows: 1) Control group (continuation of standard chow diet); 2) Western-type diet without fiber (BioServ, S3282; 60% kcal fat; irradiated; per weight: 36% fat, 20.5% protein, 35.7% carbohydrates, 0% fiber); or 3) infection with 5×10^7^ CFU of attenuated *Salmonella* Typhimurium (Stm SL1344^ΔSPI-2^). Fecal pellets were collected pre-challenge (day 0) and during the following three day. On the day 3, mice were euthanized and cecal content was collected. Fecal pellets were weighted and homogenized in 500µLof PBS as described before. Serial dilutions were prepared and plated in BHI-blood agar plates supplemented with corresponding antibiotics for CFU quantification. In addition, 100-300µL of homogenized content was plated directly for further assessment of the distribution of *B.theta*-EryR tagged strains by qPCR. Cecal content was processed as previously described.

### Quantitative PCR

Colonies from enrichment plates were pooled in 5ml of PBS and homogenized by vortex. Genomic DNA was isolated with the QIAamp DNA Mini Kit (Qiagen). qPCR was performed using with FastStart Universal SYBR Green Master Mix (Roche, Cat. N° 4385610). Primers (Supplementary Table 2) were mixed to a final concentration of 1µM. Between 160-200ng of DNA was amplified using StepOne Plus or QuantStudio 7 Flex instruments (Applied Biosystems) using the following protocol: initial denaturation at 95°C for 14min followed by 40 cycles of 94°C for 15sec, 61°C for 30sec, and 72°C for 20sec as described previously.

### Mathematical modelling

#### 1. Estimation of colonization probability based on lost tags

Let us denote *C* the bacterial concentration in the prepared solution. Then, if there is a volume *V* of solution, then there are *N* = *CV* bacteria. Therefore, the probability to have taken *n_0_* bacteria in a volume *v_0_* is:

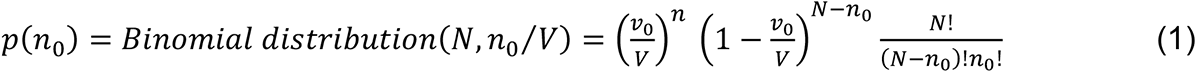

In the limit of *N* = *cV* large and *v*_0_ ≪ *V*,

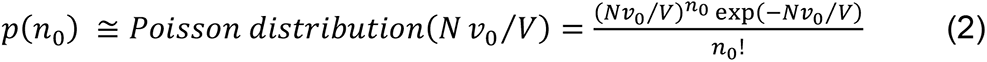

Let us define *β* as the probability for each bacterium to get to the cecum alive, and then have its lineage survive until measurement. There are a priori no interactions early on between incoming bacteria, as their concentration is initially low enough to limit the competition between them before arriving to the cecum. Then, the probability for a tagged *B.theta* strain not to be present at measurement time, if started with an average of *n_0_* bacteria (Poisson distributed) is the zero of the Poisson distribution of average *βn_0_*, and thus:

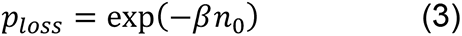

As *n_0_* is estimated via the concentration and volume of the inoculum, and *p_loss_* is best estimated via the number of tags lost divided by the total number of tags, then *β* is estimated as:

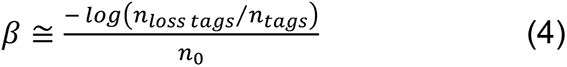

To consider the fact that not all tags have the same *n_0_*, *β* is actually estimated by its value maximizing the probability of the experimental observations, thus *β* maximizing:

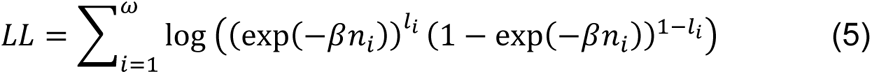

This expression is also used for calculating the confidence interval, as detailed in appendix.

#### 2. Estimation of colonization probability based on variance

The variance on the proportions is:

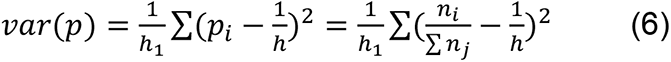

In the limit where the initial number of bacteria are of the same order of magnitude, we find:

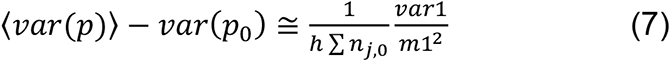

with *var*(*p*_0_) the variance in proportions in the inoculum, ∑ *n*_*j*,0_ the total number of tags in the inoculum, and *var*1/*m*1^2^ the relative variance starting from one bacterium. We find (see Supplementary Methods) that *var*1/*m*1^2^ is 2/(colonization probability). Then *var*1/*m*1^2^ is estimated for each mouse using equation (7), and the average variance is used to estimate *var*1/*m*1^2^. The standard error on *var*1/*m*1^2^ is used to obtain the confidence interval for the colonization probability.

#### 3. Estimation of clearance rate due to flow

We examined the expected magnitude of the effect of an extended lag phase in the cecum on colonization probability to determine whether this is consistent with our observed neutral tagging data. It should also be noted that the cecum is a dynamic environment with pulsatile arrival of material from the small intestine and loss of material to the feces. This generates a clearance rate due to flow, on top of any clearance rate due to bacterial death. Assuming that the main site of growth of *B.theta* is the cecum/upper colon, the parameter for clearance due to flow can be estimated by quantifying the volume of cecum content lost per day. This can be empirically estimated by measuring 1) fecal dry mass produced per day, and 2) the water content of cecum content. Assuming minimal change in dry mass during colon transit in the mice, this infers a dilution rate of cecal content in the order of 0.12 volumes/h in a germ-free mouse and 0.18 volumes/h in an SPF mouse; gnotobiotic mice will have a value in between these two. Bacterial clearance due to killing will contribute over and above these values. Of note, bacteria with a long lag-phase after introduction into the cecum will be cleared by the flow before growth can start, i.e., during the early phase of colonization this will be a determinant of colonization probability.

Estimation of cecum turnover rates

|  | Water fraction in | Dry fecal excretion (gram/day) | Estimated wet cecal mass | Wet cecal mass (gram) | Estimated cecum | Estimated cecum |
| --- | --- | --- | --- | --- | --- | --- |

|  | cecal mass<br>(%) |  | excretion<br>(gram/day) |  | turnover rate<br>(volume/day) | turnover rate<br>(volume/h) |
| --- | --- | --- | --- | --- | --- | --- |
| Germ-free | 80.9 (0.4) | 1.55 (0.27) | 8.12 (1.42) | 2.83 (0.59) | 2.87 (0.78) | 0.12 (0.03) |
| SPF | 76.2 (1.2) | 0.81 (0.09) | 3.40 (0.42) | 0.77 (0.32) | 4.42 (1.91) | 0.18 (0.08) |

#### 4. Estimation of the competitive index

We assume that bacteria have first a probability of survival *q_i_* (with *i* = *w* for the WT strain, and *i = a* for the acapsular strain). Then once the cecum is reached, they have a loss rate *c_i_*.

During an initial lag-phase *τ*_*i*_, they do not grow. Then, each bacterial strain grows logistically, initially at a rate *r*_*i*_, which saturates when approaching carrying capacity with a factor (1 - (*A* + *W*)/*K*). When carrying capacity is reached, the total number of bacteria remains constant until the end of the experiment at time *t*_*tot*_, with both loss and replication ongoing and compensating each other. Given the growth rates for WT and acapsular are similar in vitro, we assume *r*_*w*_ = *r*_*a*_, so that the difference in the initial net growth rates (*r*_*i*_ - *c*_*i*_) originates from *c*_*a*_ > *c*_*w*_. Therefore, in the competition setting, the population is mostly composed of WT when carrying capacity is reached, so that the global population size is depleted with rate approximately *c*_*w*_. As a result, the effective residual replication rate of both types exactly compensates the loss rate, *c*_*w*_, because they compete for the same resources and have the same growth ability. As acapsular keep being lost at a larger rate ca, their net growth rate at carrying capacity is negative, so that they keep decreasing in frequency in the population.

With this model (see detailed calculations in Supplementary Methods), we find that the relative ratio between WT and acapsular is:

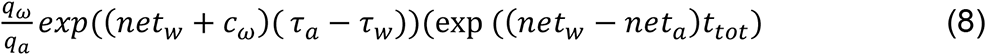

For all the microbiota except SPF, as the colonization probabilities were similar for the WT and acapsular, then we assume *q_*a*_* = *q_*w*_*. For SPF, we use the ratio of *q_*w*_* / *q_a_* in the colonization experiments, adjusted for the fact that the full colonization probability also includes steps after the initial death before reaching the cecum.

All the parameters used are determined using experiments other than the competition experiment (except for *t_tot_*, the total experimental time).

#### 5. Estimation of colonization probability during competition

We find that the overall survival probability for the acapsular in the competition experiment is the colonization probability from the colonization experiment (alone), multiplied by a factor considering later loss (when the carrying capacity is reached by the WT and the acapsular decreases). The complete expression can be found in the corresponding section of the Supplementary Methods.

#### 6. Estimation of survival probability after challenge

In these experiments, at the time of the start of the challenge, the bacterial population is at carrying capacity, so the effective growth rate (likely limited by availability of nutrients) is about the same as the loss rate. Also, let us assume that the population the population size is known at this time. Then, a challenge is applied. A challenge may have different effects: it may impose a temporary bottleneck in the population (loss becomes higher than reproduction), or it may increase the loss rate (with the reproduction rate increasing enough to compensate), and thus the turnover of the population. In any case, we can calculate *β* as the probability that a bacteria present at day=0 of the challenge has its lineage still alive at day 3. Then, if there are *n_0_* bacteria of a given type at day=0, then:

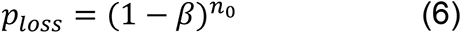

To estimate the total population size in the cecum at before the challenge (*n*_0_), we assume that (1) all the animals are in the control conditions at day=0, and (2) the cecum mass and bacteria concentration is the same on day=0 and day=3 in the control group. Therefore, the average ratio of the concentration of bacteria in feces relative to cecum at day=3 in the control animals is used to estimate the bacterial concentration in the cecum at day=0. Then, we estimate the proportion of each tag strain in from the total tagged-bacteria CFU counts using the relative distribution the qPCR data. In addition, *p*_*loss*_ is best estimated via the number of tags lost divided by the total number of tags.

## Supplementary Information

**Supplementary Figure 1:**
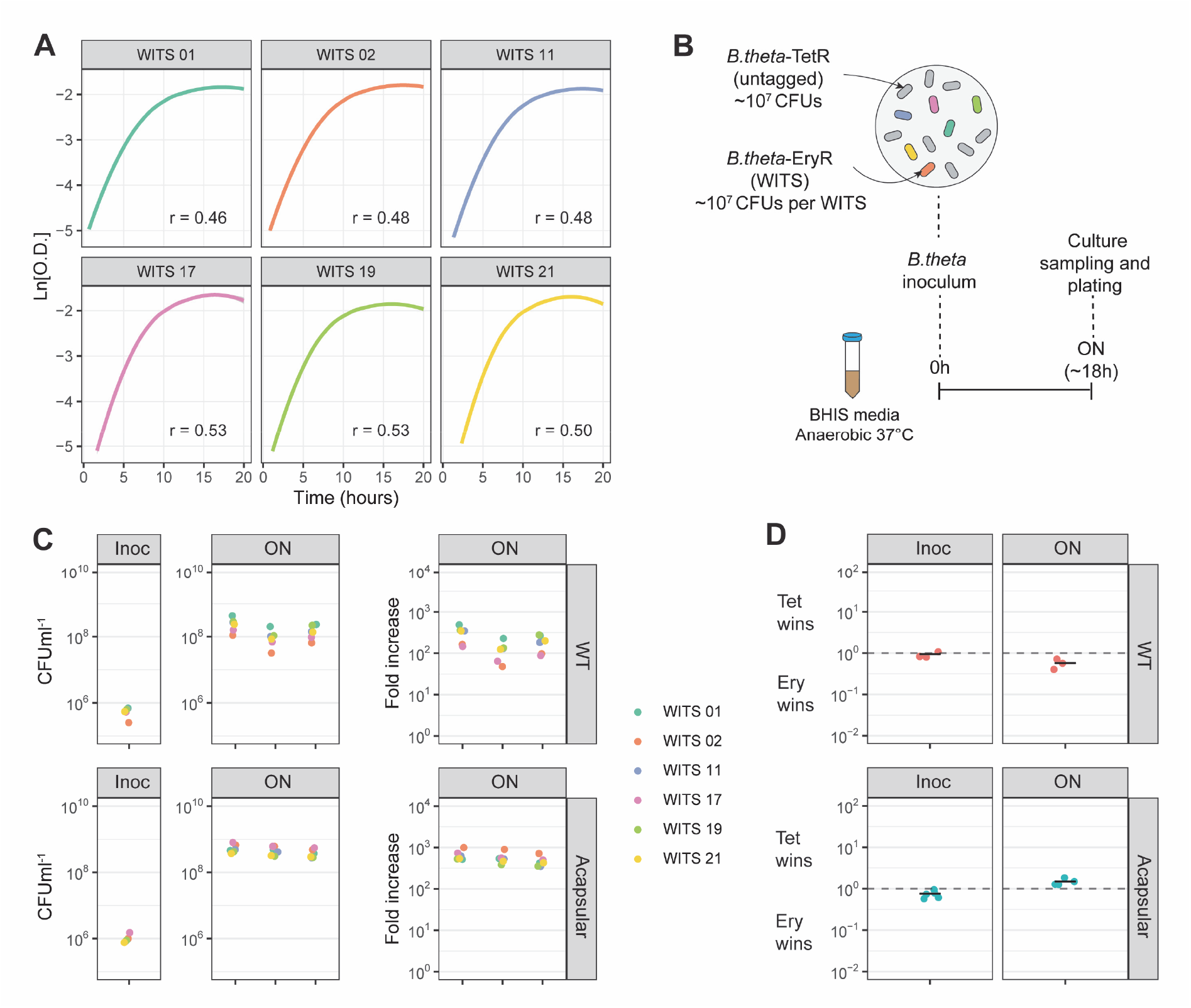
Tagged *B.theta* strains have similar fitness for growing *in vitro*. **(A)** Growth curves of *B.theta* acapsular in BHIS media and growth rates (r) per tag. **(B)** Experimental design of *in vitro* competitions. All strains (untagged *B.theta-*TetR and all six tagged *B.theta*-EryR WITS) were mixed in a 1:1 ratio. **(C)** Tags distribution after *in vitro* competition among six *B.theta* tagged strains either WT or acapsular to demonstrate neutrality *in vitro*. Plots show distribution of tags in the inoculum, in individual overnight (ON) cultures and fold increase of each tag per culture compared to the inoculum. **(D)** Competitive index of tetracycline (Tet)-over erythromycin (Ery)-resistant strains *in vitro* after ON cultures.

**Supplementary Figure 2.**
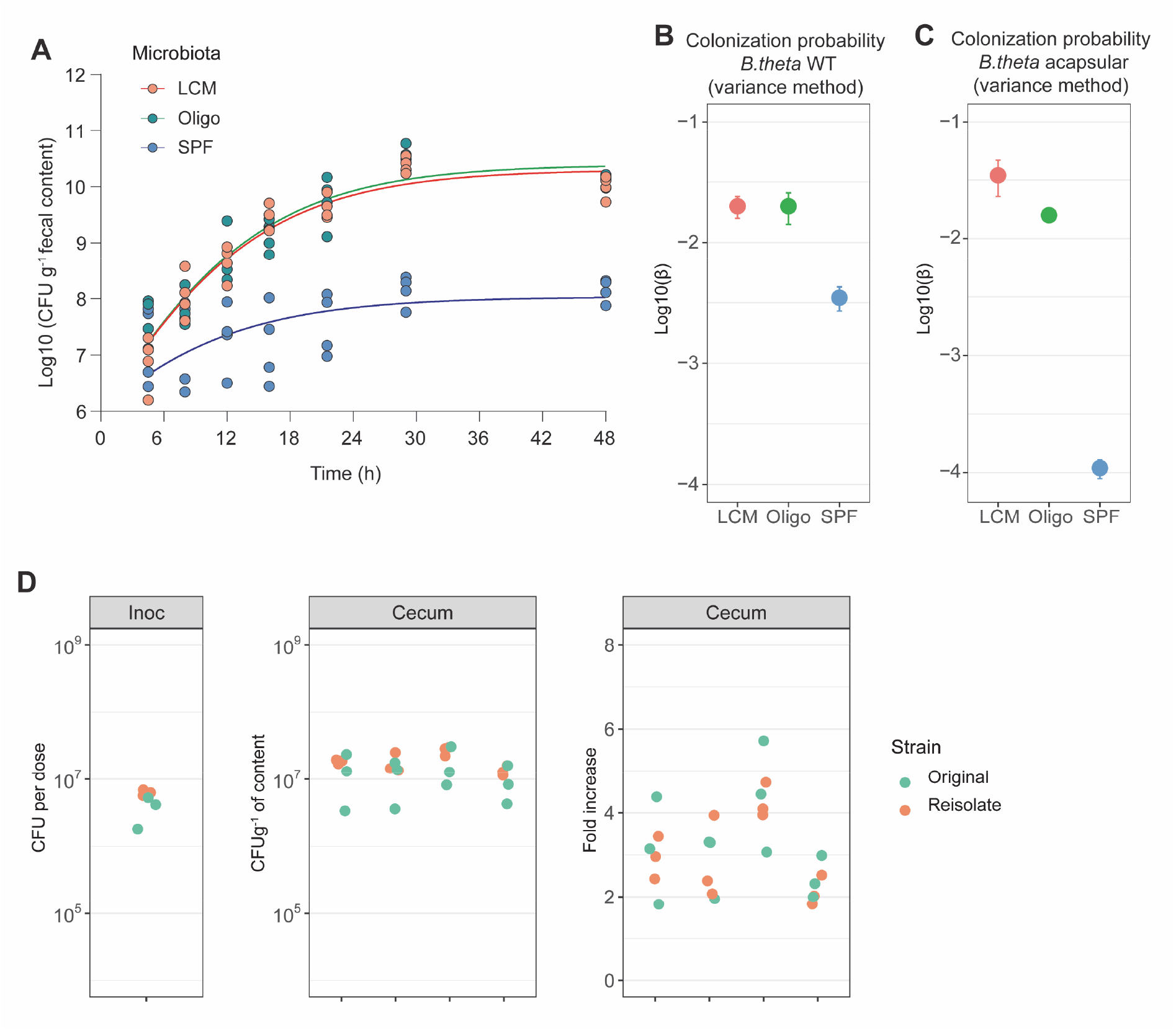
**(A)** *In vivo* growth curve of WT *B.theta* in three different microbiota backgrounds. **(B and C)** Probability of colonization (β) for **(B)** WT and **(C)** acapsular in the cecum after 48h of colonization. Estimation based on 6 tags x m mice (LCM=17, OligoMM12=10, SPF=11) and on variance on tags before (inoculum) and after colonization (cecum). **(D)** *In vivo* competition among *B.theta* strains reisolated from SPF mice vs. original stock strains. Plots show distribution of *B.theta* original and isolated strains in the inoculum, in cecal content of individual mice after 48h of colonization and fold increase of each tag per culture compared to the inoculum.

**Supplementary Figure 3.**
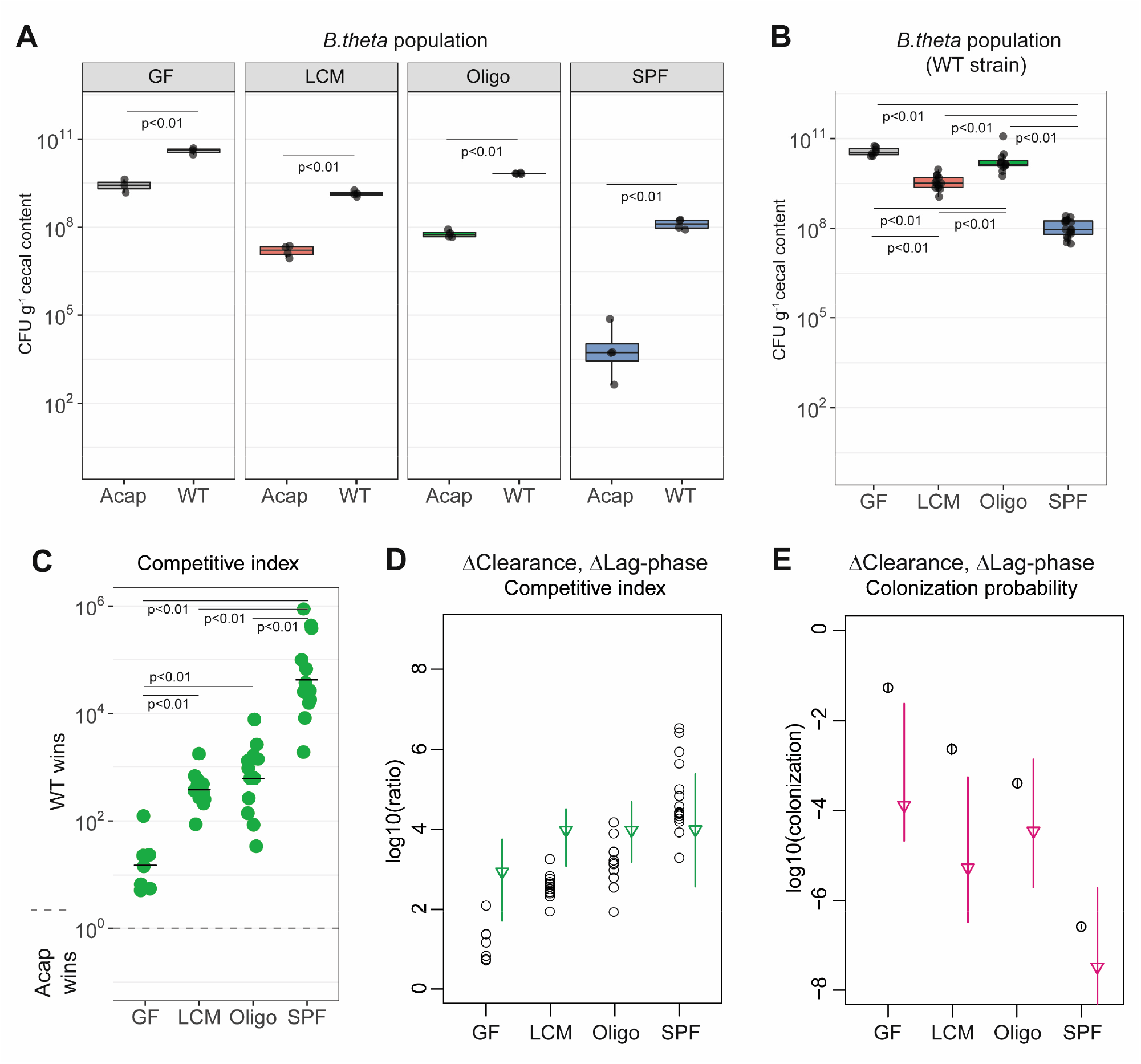
**(A)** *C*ompetition with acapsular and WT strains during colonization starting at a 1:1 ratio. Population density of the WT and acapsular strain in cecum after 48h of colonization. **(B and C)** Estimation of probability of colonization of the cecum by the acapsular strain using a WT strain inoculum spiked with tagged acapsular strains. **(B)** Population density of the WT strain and **(C)** Competitive index after correction with the initial WT/acapsular ratio in the inoculum. **(D and E)** Estimation of the **(D)** competitive index and **(E)** colonization probability of the acapsular strain assuming a mean 6h difference in lag-phase and the estimated difference in clearance rate between the WT and acapsular strains (circles: experimental data; triangles: mean parameter estimation and error).

**Supplementary Table 1:**
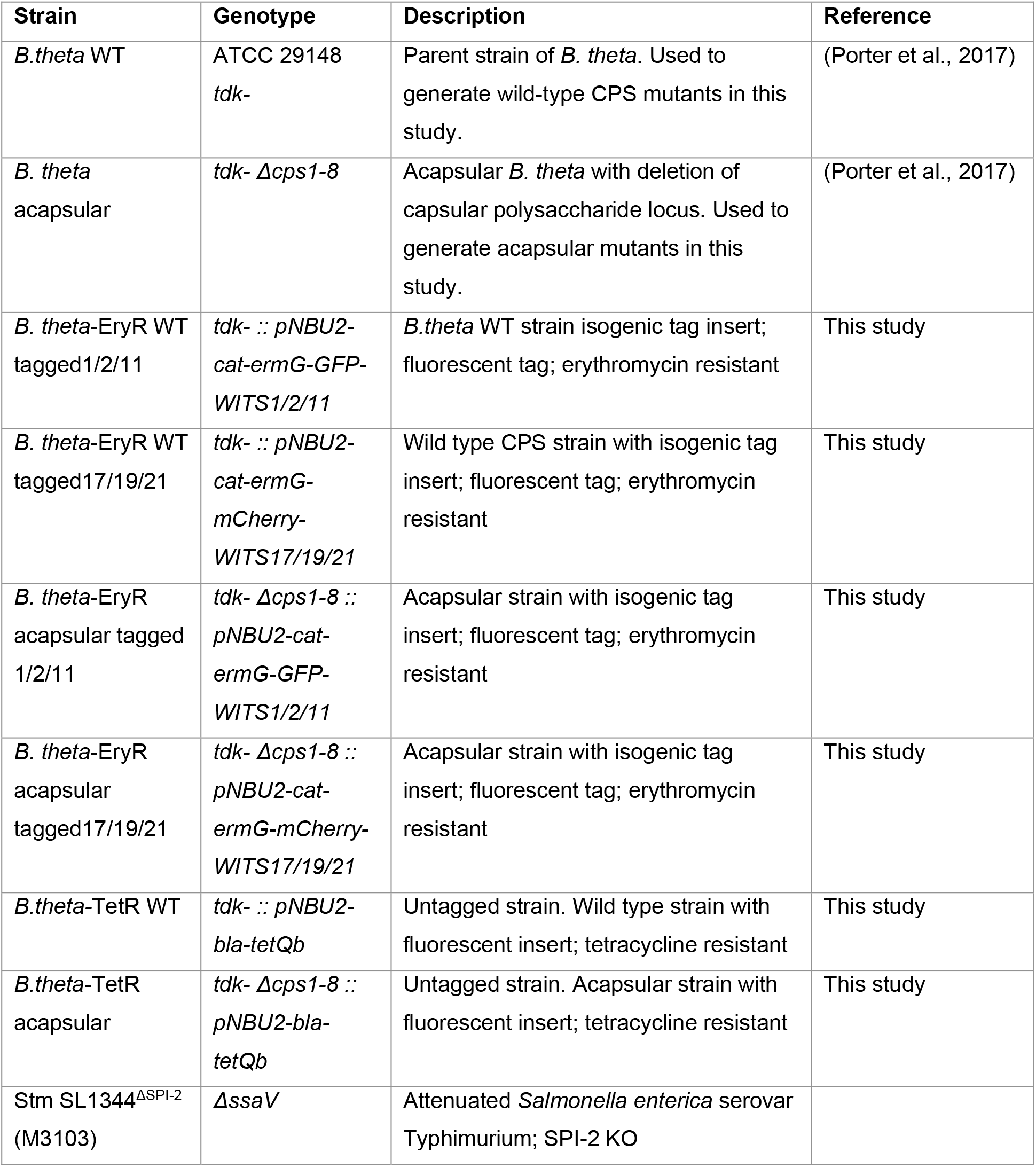
Bacterial strains

**Supplementary Table 2:**
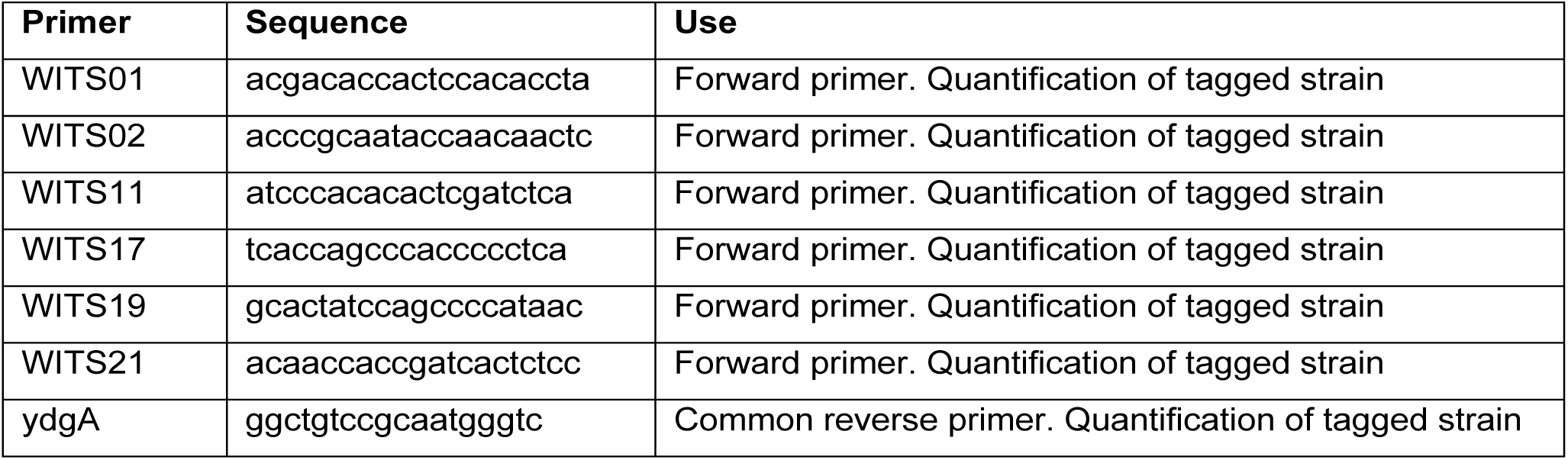
Primers

**Supplementary Table 3:** Microbiota composition in gnotobiotic mice LCM, OligoMM-12

Low complexity microbiota
| OTU | Relative abundance | 8 ASF species match | SILVA best match |
| --- | --- | --- | --- |
| 1 | 6,43E-01 | Bacteroides sp. ASF519 | Parabacteroides goldsteinii dnLKV18 |
| 2 | 1,65E-01 | Clostridium sp. ASF502 | Clostridiales bacterium |
| 3 | 9,60E-02 | Eubacterium plexicaudatum strain ASF 492 | Lachnospiraceae bacterium 6_1_63FAA |
| 4 | 5,56E-02 | Eubacterium plexicaudatum strain ASF 492 | Blautia |
| 5 | 2,00E-02 | * | Turicibacter sp. LA61 |
| 6 | 8,52E-03 | Eubacterium plexicaudatum strain ASF 492 | Lachnoclostridium;bacterium NLAE-zl-411 |
| 7 | 7,40E-03 | Bacterium ASF500 | Flavonifractor;human gut metagenome |

OligoMM12
| OTU | Strain ID | DSM number | Phyla | Classification |
| --- | --- | --- | --- | --- |
| 1 | YL2 | 26074 | Actinobacteria | *Bifidobacterium longum subsp. animalis* |
| 2 | YL27 | 28989 | Bacteroidetes | 'Muribaculum intestinale' |
| 3 | I48 | 26085 | Bacteroidetes | 'Bacteroides caecimuris' |
| 4 | YL45 | 26109 | Proteobacteria | 'Turicimonas muris' |
| 5 | YL44 | 26127 | Verrucomicrobia | *Akkermansia muciniphila* |
| 6 | KB1 | 32036 | Firmicutes | *Enterococcus faecalis* |
| 7 | I49 | 32035 | Firmicutes | *Lactobacillus reuteri* |
| 8 | YL32 | 26114 | Firmicutes | *Clostridium clostridioforme* |
| 9 | YL58 | 26115 | Firmicutes | *Blautia coccoides* |
| 10 | YL31 | 26117 | Firmicutes | *Flavonifractor plautii* |
| 11 | KB18 | 26090 | Firmicutes | 'Acutalibacter muris' |
| 12 | I46 | 26113 | Firmicutes | *Clostridium innocuum* |

## Notes

### Competing Interest Statement

The authors have declared no competing interest.

