## Supplementary Methods for "Fitness advantage of *Bacteroides thetaiotaomicron* capsular polysaccharide is dependent on the resident microbiota"

2.3 Colonization probability via the loss - different starting inoculum sizes - best estimate . . 6
2.4 Colonization probability via the loss - different starting inoculum sizes - confidence interval 6

### 1 Optimal number of tagged bacteria spiked in the inoculum ($n_0$)

#### 1.1 Principle

Here $\beta$ is the probability for the lineage of one bacterium, present in the inoculum, to be found at measurement time. It is a combination of arriving alive to the cecum, and not being lost afterwards. $N$ is the total number of tagged bacteria measured (the number of tagged bacteria in each mouse multiplied by the number of mice).

There are two main options when computing the estimated $\beta$:

- Either take $\beta$ as 0 when all the tags are lost, and take $\beta$ as 1 when no tags are lost.
- Or do not try to estimate $\beta$ in both these situations.

#### 1.2 Analytical results

The probability for the observation of $n_l$ lost tags is

$$p(n_l) = \exp(-\beta n_0)^{n_l}(1 - \exp(-\beta n_0))^{N-n_l}\frac{N!}{(N-n_l)!n_l!} \tag{1}$$

##### 1.2.1 Probability of all lost and none lost

The probability to observe the loss of all tags is:

$$p(\textit{all lost}) = \exp(-\beta n_0 N) \tag{2}$$

The probability to observe no tags loss is:

$$p(\textit{no loss}) = (1 - \exp(-\beta n_0))^N \tag{3}$$

##### 1.2.2 Mean estimated $\beta$

It can be checked that, provided that the probability that having all or none tags lost is small, the average estimated $\beta$ corresponds to the true $\beta$.

##### 1.2.3 Variance on the estimate of $\beta$

The next step is to look where the typical relative error on the estimate is the smallest, i.e. where the relative variance is minimized:

In the case for which we always estimate $\beta$,

$$\frac{var}{\beta^2} = (\frac{1}{\beta}-1)^2(1-\exp(-\beta n_0))^N+\exp(-\beta n_0 N)+\sum_{n_l=1}^{N-1}\frac{(-\log(n_l/N)-\beta)^2}{\beta^2 n_0}\exp(-\beta n_0)^{n_l}(1-\exp(-\beta n_0))^{N-n_l}\frac{N!}{(N-n_l)!n_l!} \tag{4}$$

In the case for which we do not estimate $\beta$ when either all or no tags are lost:

$$\frac{var}{\beta^2} = \frac{\sum_{n_l=1}^{N-1}\frac{(-\log(n_l/N)-\beta)^2}{n_0\beta^2}\exp(-\beta n_0)^{n_l}(1-\exp(-\beta n_0))^{N-n_l}\frac{N!}{(N-n_l)!n_l!}}{1-(1-\exp(-\beta n_0))^N - \exp(-\beta n_0 N)} \tag{5}$$

These expressions are not easily analytically solvable. Let us place ourselves in the case when $\beta n_0$ is neither small ($\gg 1/N$) nor large ($\ll N$). In this regime, loosing all or no tags is unlikely, so both expressions are equivalent. We take the largest terms, defining $\epsilon$ as $n_l \simeq (1+\epsilon)N\exp(-\beta n_0)$, and obtain:

$$\frac{var}{\beta^2} \simeq \sum_{n_\epsilon}\frac{(-\log(1+\epsilon)/\beta + n_0 - 1)^2}{n_0}\exp(-\beta n_0)^{n_l}(1-\exp(-\beta n_0))^{N-n_l}\frac{N!}{(N-n_l)!n_l!} \tag{6}$$

Assuming $\log(1+\epsilon) \simeq \epsilon = n_l\exp(\beta n_0)/N - 1$

$$\frac{var}{\beta^2} \simeq \sum_{n_l}\frac{(-n_l\exp(\beta n_0)/(N\beta) - 1/\beta + n_0 - 1)^2}{n_0}\exp(-\beta n_0)^{n_l}(1-\exp(-\beta n_0))^{N-n_l}\frac{N!}{(N-n_l)!n_l!} \tag{7}$$

$$\frac{var}{\beta^2} \simeq \frac{1+\exp(-\beta n_0)(-1+N(\beta(n_0-1)-2)^2))}{\beta^2 n_0 \exp(-\beta n_0) N} \tag{8}$$

Assuming that $N$ is large, the expression is minimized for $\beta(n_0-1)-2 \simeq 0$, i.e. $n_0 \simeq 2/\beta$.

As there are assumptions to obtain this result, we further confirm it numerically (next section).

#### 1.3 Numerical results

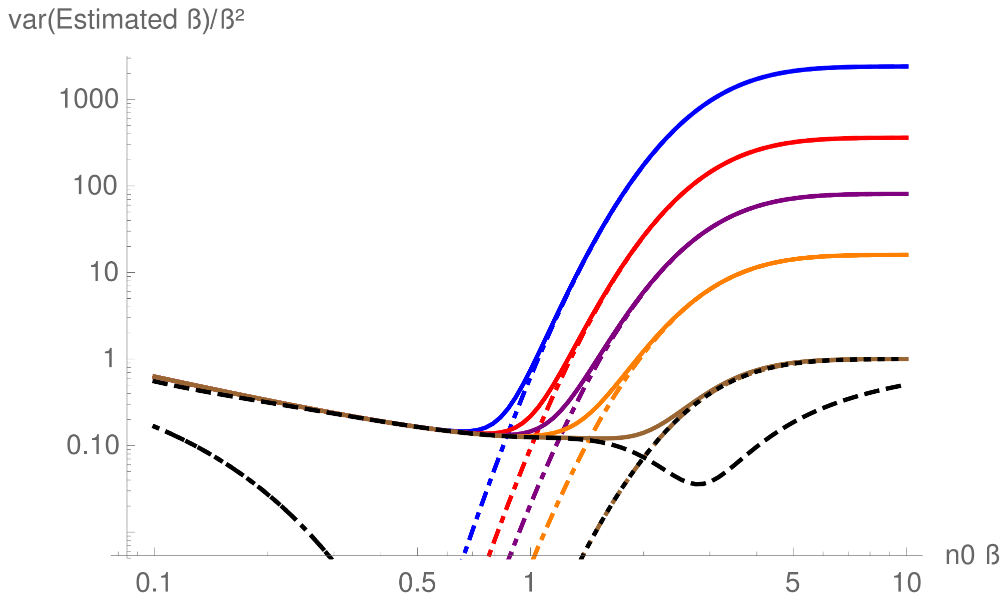

Figure 1: The relative variance in the estimate of $\beta$, as a function of $n_0\beta$. The solid colored lines are the expression (4) for $\beta = 0.02, 0.05, 0.1, 0.2, 0.5$. The black dashed line the expression (5) (which depends on $\beta n_0$ but not on $\beta$ separetely, thus is the same here for any $\beta$). The coloured dot-dashed lines are $(\frac{1}{\beta}-1)^2(1-\exp(-\beta n_0))^N$, the term due to the estimate of $\beta$ as 1 when no tags are lost, and the black dot-dashed line is $\exp(-\beta n_0 N)$, the term due to the estimate of $\beta$ as 0 when all tags are lost. The black dotted line represents the probability to be in one of these cases. $N = 18$.

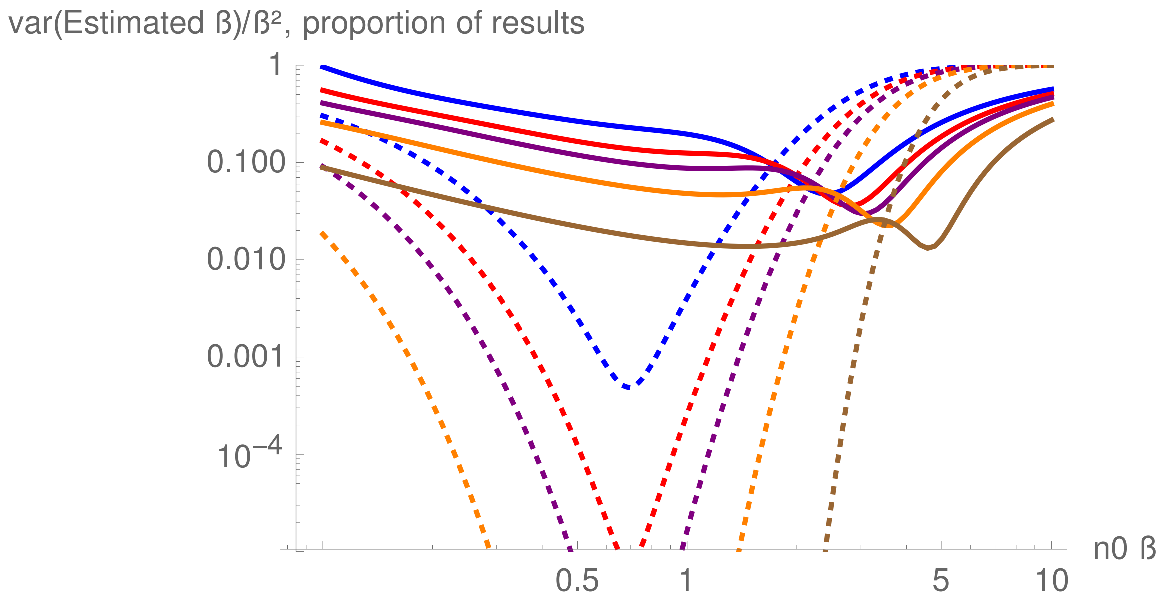

Figure 2: The relative variance in the estimate of $\beta$, as a function of $n_0\beta$. The solid colored lines are the expression (5) for $N = 12, 18, 24, 40, 120$. The dotted lines is the probability that of either all or no tags are loss.

Figure 1 and 2: if $\beta$ is taken as 1 when no tags are lost, there is a risk of overestimating it, in particular for large $\beta n_0$. When looking at the more reasonable case where $\beta$ is estimated only if there is at least one tag lost (and at least one tags present), while there seems to be a minimum of the relative variance for $\beta n_0$ of several units, increasing with $N$, this minimum is for a relatively high probability of not producing any estimate. Then, the minimum of the relative variance with high probability of providing an estimate is when $\beta n_0$ is in the order of 1. Therefore, to ensure that some tag loss is always observed when $N$ is relatively small (under 50), then $\beta n_0$ should be just below 1.

#### 1.4 Conclusion

The value of $n_0$ leading to the best $\beta$ estimates (likely possibility of obtaining an estimate, accurate average, minimized relative variance between the estimate and the true value) is such that $\beta n_0$ is of the order of 1, and when $N$ (the number of tagged bacteria per mouse multiplied by the number of mice) is small, it is safer to have it a bit smaller.

#### 2 Colonization

##### 2.1 Initial bacteria number distribution

Let us denote $c$ as the bacterial concentration in the prepared solution. Then, if there is a volume $V$ of solution, there are $N = cV$ bacteria. Then, the probability to have taken $n_0$ bacteria in the volume $v_0$ of the inoculum is:

$$p(n_0) = Binomial\ Distribution(N, v_0/V) = \left(\frac{v_0}{V}\right)^{n_0}\left(1-\frac{v_0}{V}\right)^{N-n_0}\frac{N!}{(N-n_0)!n_0!} \tag{9}$$

In the limit of $N = cV$ large and $v_0 \ll V$,

$$p(n_0) \simeq Poisson\ Distribution(Nv_0/V) = \frac{\left(\frac{Nv_0}{V}\right)^{n_0}\exp\left(-\frac{Nv_0}{V}\right)}{n_0!} \tag{10}$$

##### 2.2 Colonization probability via the loss - base theory

Let us define $\beta$ as the probability for each bacterium clone to get to the cecum alive, and then have its lineage survive until measurement. There is *a priori* no interaction early on between incoming bacteria, as their concentration is initially low enough to limit the competition between them. Then, the probability for a tagged bacterium to not be present at measurement time, if started with an average of $n_0$ bacteria (Poisson distributed) is the zero of the Poisson distribution of average $\beta n_0$, and thus:

$$p_{loss} = \exp(-\beta n_0). \tag{11}$$

Then, as $n_0$ is estimated via the concentration and volume of the inoculum, and $p_{loss}$ is best estimated via the number of tags lost divided by the total number of tags, $\beta$ is estimated as:

$$\beta \simeq \frac{-\log(n_{lost\ tags}/n_{tags})}{n_0}. \tag{12}$$

##### 2.3 Colonization probability via the loss - different starting inoculum sizes - best estimate

The tagged bacteria are not necessarily in equal concentrations in the inoculum and the data from several experiments with different inoculums need to be combined. Let us define $w$ the number of tags multiplied by the number of mice. For each of these $w$, there were $n_i$ tagged bacteria in the inoculum, and we define $l_i$ as 1 if the tag was lost, 0 otherwise. Then, the estimate of $\beta$ is $\beta$, which maximizes $proba(l_1, ...., l_w)$ the likelihood to observe $\{l_1, l_2, ...l_w\}$; and it is the same as maximizing the log likelihood. As for each tagged bacterium the process will be considered as independent, it will be simply the maximization of:

$$LL = \sum_{i=1}^{w}\log\left((\exp(-\beta n_i))^{l_i}(1-\exp(-\beta n_i))^{1-l_i}\right) \tag{13}$$

This is the same as maximizing the following expression

$$LL' = \sum_{i=1}^{w}(-l_i\beta n_i + (1-l_i)\log(1-\exp(-\beta n_i))) \tag{14}$$

Let's note $x = -\beta$.

$$\frac{dLL'}{dx} = \sum_{i=1}^{w} n_i\left(l_i - (1-l_i)\frac{\exp(xn_i)}{1-\exp(xn_i)}\right) \tag{15}$$

The value of $x$ that makes this expression zero is found numerically, and enables to infer $\beta$, and enable to combine data with different initial number of bacteria in the inoculum coming from different mice.

##### 2.4 Colonization probability via the loss - different starting inoculum sizes - confidence interval

To calculate a confidence interval, it is useful to obtain an estimate of the probability that a given $\beta$ is the true value knowing the given observations, which can be denoted $p(\beta|observations)$. Therefore, we used a Bayesian approach, $p(\beta|observations) = p(observations|\beta)p(\beta)/p(observations)$. In this expression, $p(observations|\beta)$ is the probability to observe $\{l_1, l_2, ...l_w\}$ for a given $\beta$, so it is actually $\exp(LL)$. Then, the prior on $p(\beta)$ has to be chosen. As it may be frequent to have a very low probability of survival, a prior flat for $p(\log(\beta))$ may be better than a prior flat for $p(\beta)$. Then a similar reasoning can be done, $p(\log(\beta)|observations) = p(observations|\log(\beta))p(\log(\beta))/p(observations)$, and with a flat prior for $p(\log(\beta))$, $p(\log(\beta|observations)) \propto p(observations|\log(\beta))$. Then, $\exp(LL(\log(\beta))$ renormalized by its integral for $\log(\beta)$ gives an estimate of the distribution of probability of inference of $\log(\beta)$. The exponential of the average of $\log(\beta)$ on this distribution is very close to the value of $\beta$ maximizing $LL'$. This distribution is very close to a Gaussian distribution (see Figure 3). Then the mean $\log(\beta) \pm$ twice the standard deviation of $\log(\beta)$ on this distribution gives a 95% confidence interval.

Note that if we had chosen a prior flat on $p(\beta)$ rather than $p(\log(\beta))$, the result would have been very similar, though with a mean slightly further away from $\beta$ maximizing $LL'$, and the distribution would be less similar to a Normal distribution.

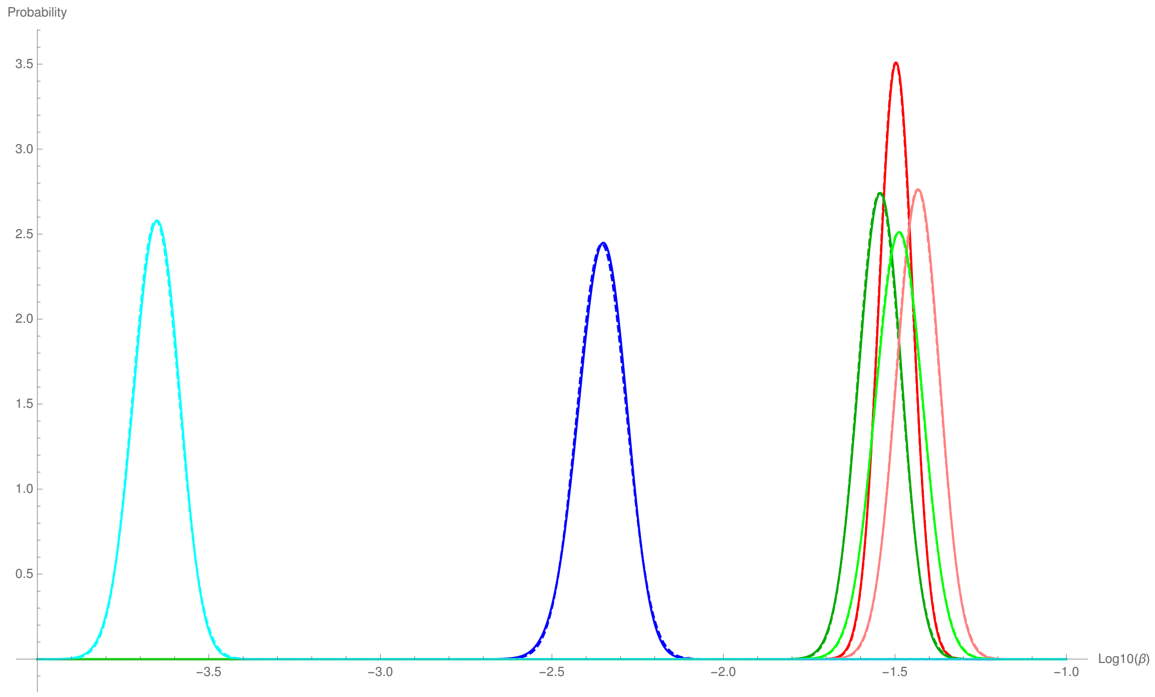

Figure 3: Distribution of the probability estimation of $\beta$, as a function of $\log_{10}(\beta)$. The colors represent experiments: LCM WT, Oligo WT, SPF WT, LCM acapsular, Oligo acapsular, SPF acapsular. Solid lines represent the numerical result for the probability distribution of $\beta$, using renormalized equation (14), the dashed lines represent the Normal distribution for $\log_{10}(\beta)$, with the same mean $\log_{10}(\beta)$ and same variance than the numerical distribution.

#### 2.5 Colonization probability via the loss - results

The results here are the values of $\beta$ maximizing the log likelihood, with the confidence interval as explained in previous section:

| | LCM | Oligo | SPF |
|---|---|---|---|
| WT | $10^{-1.50[-1.60:-1.40]}$ | $10^{-1.54[-1.67:-1.42]}$ | $10^{-2.35[-2.50:-2.21]}$ |
| Acapsular | $10^{-1.43[-1.56:-1.31]}$ | $10^{-1.49[-1.63:-1.35]}$ | $10^{-3.65[-3.79:-3.52]}$ |

In principle, as we make multiple comparisons, the confidence interval should be adjusted, but here, even without correction, the LCM/Oligo mice for both WT and acapsular are not significantly different (adjusting for multiple comparisons would make them even less distinguishable). In the case of SPF mice, the WT strain is more than 10 standard deviations away for the WT bacteria, and more than 30 standard deviations for the acapsular bacteria, relative to LCM/Oligo mice. Within the SPF mice, the difference between WT and acapsular bacteria is more than 18 standard deviations. Because the differences are very large, even when adjusting for multiple comparisons, they would remain significant.

#### 2.6 Colonization probability via the loss - Interpretation of the apparent loss probability: initial loss and subsequent loss

##### 2.6.1 Principle

What is obtained here is the probability for a bacterium clone in the inoculum to have seeded a lineage still present at measurement. For the lineage to be still present, it needs to:

- make it alive to the cecum. Let us denote this probability $q$.
- escape stochastic fluctuations, mainly during the initial rounds of reproduction.

If there is little subsequent loss, then $\beta \simeq q$.

##### 2.6.2 Minimal model

Here, we assume that each bacterium has a probability $q$ of establishing, and then its lineage grows at a rate $r$ and is lost at a rate $c$. After some time, carrying capacity is reached. Then, we assume that the population is large, and thus stochastic lost will be negligible at this point. Therefore, as the stochastic effects occur when the population size is small, then it is legitimate to focus on this step.

The corresponding generating function for the population size at time $t$ is:

$$g(z,t) = 1 - q\frac{(r-c)(1-z)}{(rz-c)\exp(-(r-c)t) + r(1-z)} \tag{16}$$

The resulting loss probability is:

$$p_{loss} = g(0,t) = 1 - q\frac{r-c}{r - c\exp(-(r-c)t)} \xrightarrow{(r-c)t \gg 1} 1 - q\left(1 - \frac{c}{r}\right) \tag{17}$$

##### 2.6.3 Model with delay

As shown in Fig. 3E, we observed a delay in the start of growth in the mouse gut, during which time bacteria will continue to be lost due to flow, but will not replicate. Therefore, in a model with a fixed delay, it would be the same, except with $q\exp(-c\tau)$ instead of $q$.

#### 2.7 Colonization - Using the variance in the size of the different tagged population - theory

##### 2.7.1 Simple bottleneck

Here, we assume that each bacterium has a probability $\beta$ of establishing.

Then:

$$p_{loss} = 1 - \beta \tag{18}$$

And the variance for $\beta$:

$$var = \beta(1-\beta)^2 + (1-\beta)\beta^2 = \beta(1-\beta) \tag{19}$$

$$var_{rel} = \frac{var}{\beta^2} = \frac{1-\beta}{\beta} \xrightarrow{\beta \ll 1} \frac{1}{\beta} \tag{20}$$

Here $var_{rel} = 1/(1 - p_{loss})$.

##### 2.7.2 Bottleneck, replication rate and loss rate

Similar the previously described minimal model, we assume that each bacterium has a probability $q$ of establishing, and then its lineage grows at a rate $r$ and is lost at a rate $c$.

As seen in the previous section, the corresponding generating function for the population size at time $t$ is:

$$g(z,t) = 1 - q\frac{(r-c)(1-z)}{(rz-c)\exp(-(r-c)t) + r(1-z)} \tag{21}$$

and the resulting loss probability is:

$$p_{loss} = g(0,t) \xrightarrow{(r-c)t \gg 1} 1 - q\left(1 - \frac{c}{r}\right) \tag{22}$$

and the resulting variance for the population size is:

$$var = \left(\frac{\partial^2 g}{\partial z^2} + \frac{\partial g}{\partial z} - \left(\frac{\partial g}{\partial z}\right)^2\right)_{z=1} \tag{23}$$

$$var = q^2 e^{2(r-c)t}\left(\frac{1}{q(1-\frac{c}{r})}\left(2 - (1+\frac{c}{r})e^{-(r-c)t}\right) - \frac{1-\frac{c}{r}}{1+\frac{c}{r}}\right) \tag{24}$$

$$var \xrightarrow{(r-c)t \gg 1} q^2 e^{2(r-c)t}\left(\frac{2}{q(1-\frac{c}{r})} - \frac{1-\frac{c}{r}}{1+\frac{c}{r}}\right) \tag{25}$$

$$var_{rel} = \frac{var}{(q\exp((r-c)t))^2} = \frac{2}{q(1-c/r)} - \frac{1-c/r}{1+c/r} \xrightarrow{q \ll 1} \frac{2}{q(1-c/r)} \tag{26}$$

Here $var_{rel} = 2/(1-p_{loss})$.

##### 2.7.3 Different initial number of bacteria

In the limit where the initial number of bacteria are of the same order of magnitude, and with $w$ the number of different tags, the variance on the proportions is:

$$var(p) = \frac{1}{w-1}\sum(p_i - 1/w)^2 = \frac{1}{w-1}\sum\left(\frac{n_i}{\sum n_j} - \frac{1}{w}\right)^2 \tag{27}$$

After some calculations, we find:

$$\langle var(p)\rangle - var\ p_0 \simeq \frac{1}{w\sum n_{j,0}}\frac{var_1}{m_1^2} \tag{28}$$

with $var\ p_0$ the variance in proportions in the inoculum, $\sum n_{j,0}$ the total number of tagged bacteria in the inoculum, and $var_1/m_1^2$ the relative variance starting from one bacteria, which is approximately $\frac{2}{q(1-c/r)}$ and thus expected to be approximately equal to $2/\beta$.

Then:

$$\frac{var_1}{m_1^2} \simeq w\sum n_{j,0}\left(\langle var(p)\rangle - var\ p_0\right) \tag{29}$$

##### 2.7.4 Procedure

The procedure is then to estimate $\frac{var_1}{m_1^2}$ for each mouse using this equation, and then average the results to obtain $\langle\frac{var_1}{m_1^2}\rangle$, and

$$\beta \simeq \frac{2}{\langle var_1/m_1^2\rangle}. \tag{30}$$

There is one mouse (for the acapsular in SPF) for which all the tagged bacteria were lost. We remove this mouse from the analysis, but keep the 14 others so that the bias is likely minimal.

Then twice the standard error (the standard error is the standard deviation divided by the square root of the number of mice for each condition, reflecting that if there are more mice, a better the average is determined) is used for the confidence interval around $\langle\frac{var_1}{m_1^2}\rangle$, which bounds are then used to define the bounds of the confidence interval for the estimate of $\beta$ via the variance.

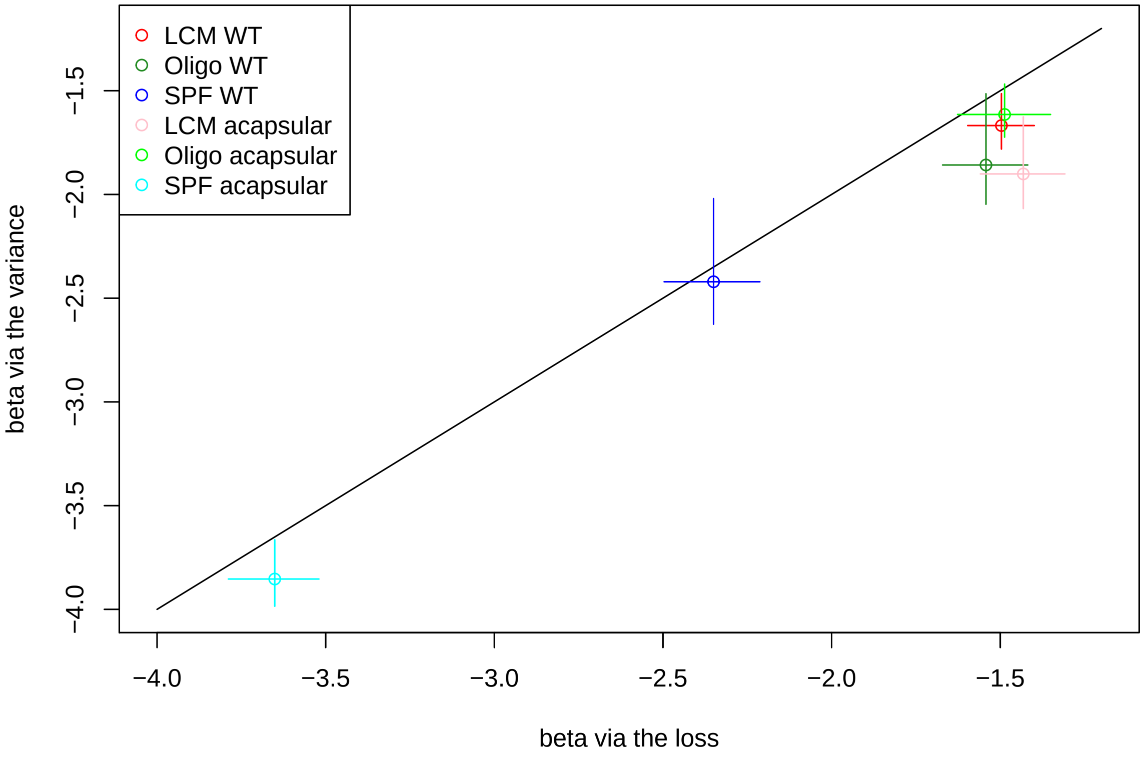

Figure 4: Comparison between the lineage survival probability estimated via the loss method, and via the variance method (circle marker and error bars). The colors represent experiments: LCM WT, Oligo WT, SPF WT, LCM acapsular, Oligo acapsular, SPF acapsular. The error bars are given via the direct study of the estimation probability for the loss method; via the standard error between mice for the variance method.

#### 2.8 Colonization - Using the variance in the tagged strain population sizes - Results and comparison with tagged strain loss

| | LCM | Oligo | SPF |
|---|---|---|---|
| WT | $10^{-1.66[-1.78:-1.51]}$ | $10^{-1.85[-2.05:-1.51]}$ | $10^{-2.42[-2.62:-2.02]}$ |
| Acapsular | $10^{-1.90[-2.07:-1.63]}$ | $10^{-1.61[-1.72:-1.47]}$ | $10^{-3.85[-3.99:-3.66]}$ |

Figure 4 compares the loss and the variance method. Overall:

- Both methods give the same orders of magnitude.
- Both methods show that in this dataset, there is no significant difference for all the experiments in LCM and Oligo mice; while WT in SPF mice has a tighter bottleneck (thought not really significant in the variance method), and acapsular in SPF mice even more.
- Overall, the variance method results in smaller $\beta$ estimate. The variance method is more sensitive on the model used, and approximations made in the minimal model may explain this systematic difference.

#### 2.9 Experimental noise

There are incertitudes on measurements:

- To estimate overall bacterial load, a number of colonies $n$ is counted, and so we expect a typical relative error of $1/\sqrt{n}$.

- The relative proportion of the different tagged strains are analyzed by qPCR. In one experiment, 3 replicates were measured for each of the tagged strain, and it showed a typical incertitude of 0.22.

For the loss method, the end point measurement is the absence/presence of a tagged strain, which is not sensitive to the qPCR incertitude (at least at the small level qPCR noise of the experiments). However, the calculations actually estimate $\beta n_i$, thus errors in the estimate of $n_i$, the initial expected number of bacteria for each tagged strain, will affect the estimate of $\beta$.

For instance, the typical number of colonies counted for checking the initial concentration from which $n_0$ is calculated is of the order of 40, resulting in a typical relative error of $1/\sqrt{40}$, i.e. about 15%. Then overestimating $n_0$ by 15% will result in underestimating $\beta$ by about 15%. The relative error is not biased, it will just increase the incertitude. $\pm 15\%$ will corresponds to approximately $\pm 0.06$ in the $log10(\beta)$. This is if all data was from the same inoculum. Actually, for each condition, the data is pooled from different experiments, with different starting inoculums, with uncorrelated incertitude on the initial number of tagged bacteria (2 to 5 different conditions depending on the condition). Then, this incertitude is smaller than the incertitude as calculated previously, thus, while it would somewhat increase the confidence interval, it has a small impact, and it is not included in our main results for simplicity.

The impact of the qPCR incertitude on the loss method is smaller. Indeed, the incertitude due to qPCR counts for one tagged strain is of the same order of magnitude as the incertitude on total $n_0$ discussed in previous paragraph; but then for each inoculum there are 6 tagged strains, and the total number of tagged bacteria is fixed, thus the overestimates and underestimates will almost compensate, and the resulting incertitude will be small compared to the one on $n_0$.

For the variance method, after some calculations, we find that the expressions can be modified to take into account the incertitude on $n_0$, and with $\sigma$ the standard deviation in noise for the number of counts,

$$\langle var_p \rangle \simeq var_{p0} + \frac{var_1}{m_1^2} \frac{1}{h^2 \langle n_0 \rangle} + \frac{1}{h^2 \langle n_0 \rangle} + \frac{2 \log(2)^2 \sigma^2}{h^2}. \tag{31}$$

With the experimental values, this correction is very small and thus is not included in the main results for simplicity.

### 3 Competition

#### 3.1 General dynamics

The WT and acapsular are thought to interact only through competition for food. After a first bottleneck of survival probability $q_i$ for each initial bacteria, their dynamics consists of:

- Fixed loss rate of $c_i$ (with $i$ standing for either WT or acapsular; and the loss rate at least equal to the cecum turnover rate)
- With a maximal carrying capacity $K_{max}$, a growth rate of $r_i(1-(W+A)/K_{max})$, with $r_i$ the maximal growth rate, $W$ the WT load, and $A$ the acapsular load.

The effective carrying capacity for the WT is $K = K_{max}(1 - c_w/r_w)$.

As depicted in Figure 5, there are few WT and acapsular bacteria early during colonization, so they grow at their maximal rate. We will approximate the more realistic logistic growth by a exponential growth until reaching the effective carrying capacity. The difference is very small and it is much easier to handle analytically. Then in the experiments we analyzed, given that they started in similar concentrations, and that the acapsular has a smaller net growth rate when the WT reaches carrying capacity, the WT hits first the carrying capacity at $t = t_w$ ($t_w$ is such that $K = n_{0w}q_w \exp((r_w - c_w)t_w)$, at which point $W \gg A$, so that $W + A \simeq W \simeq K$, at which point the net growth rate of the acapsular is:

$$net'_a = r_a(1-K/K_{max}) - c_a = r_a c_w/r_w + net_a - r_a = net_a - r_a(1-c_w/r_w) = net_a - rr net_w \quad (32)$$

which is smaller than 0 if $rr = r_a/r_w$ is close to 1, as $net_a = r_a - c_a < r_w - c_w = net_w$. The growth data showed that the net growth rate is smaller for the acapsular; and that the acapsular in competition decreases once the WT has hit carrying capacity (Fig.3F in main text).

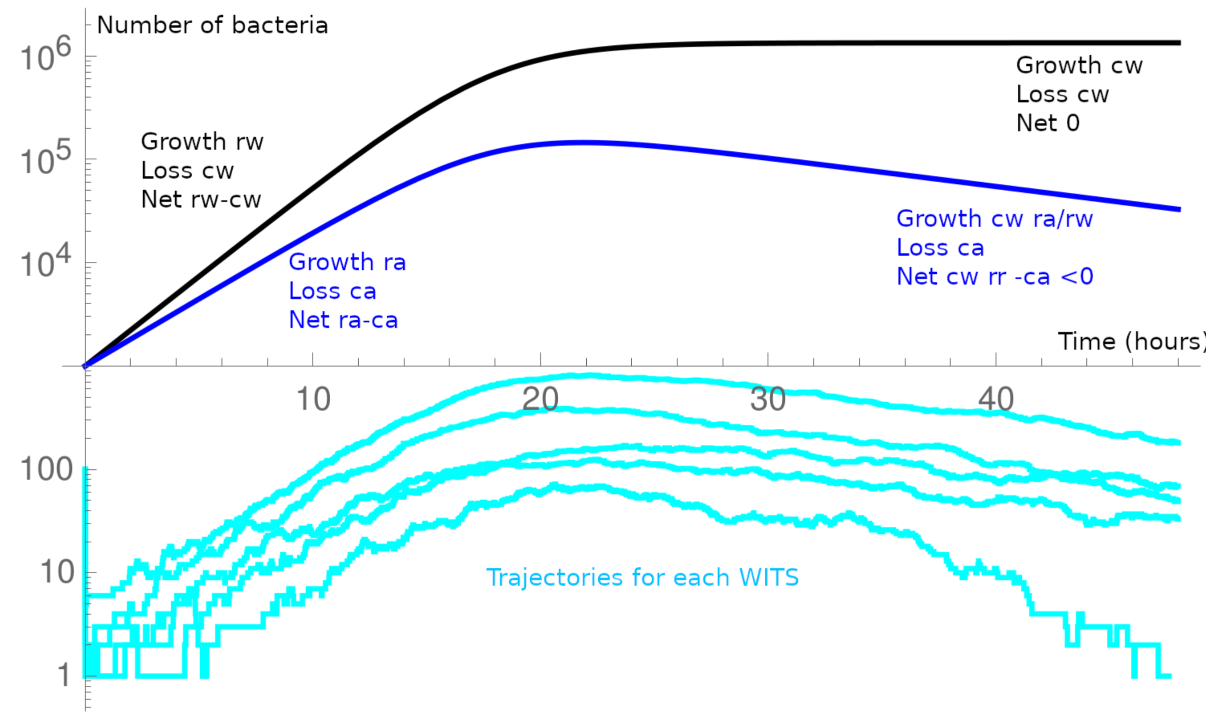

Figure 5: Schematic of the general dynamics in the minimal model. The WT (black) and acapsular (tagged strains in blue) population are in competition. After a first bottleneck (at $t = 0$), as the acapsular has a larger loss rate, its net growth rate is smaller. As the bacteria only interact through food, they follow their own dynamics until carrying capacity is reached, then WT is with a null net growth rate, and the acapsular, with its higher loss rate, has a negative net growth rate (we usually assume the absolute growth rate are equal, i.e. $rr = r_a/r_w = 1$). The tagged strains may be lost at the first bottleneck; or in the initial dynamics at low numbers; or towards the end of the experiment when the acapsular population decreases.

The generating function for the number of acapsular bacteria is $1 - q_a + q_a g_{up}(g_{plat}(z))$ with:

$$g_{up}(z) = 1 - \frac{net_a(1-z)\exp(net_a t_w)}{r_a z - c_a + (1-z) r_a \exp(net_a t_w)} \tag{33}$$

$$g_{plat}(z) = 1 - \frac{net'_a(1-z)\exp(net'_a(t_{tot} - t_w))}{r_a \frac{c_w}{r_w} z - c_a + (1-z) r_a \frac{c_w}{r_w} \exp(net'_a(t_{tot} - t_w))} \tag{34}$$

and thus the overall survival probability is $q_a(1 - g_{up}(g_{plat}(0)))$.

When bacteria are not in competition, they reach a large carrying capacity, so later loss is negligible, resulting in a total survival probability for one bacteria of:

$$q_{i,app} = q_i(1 - c_i/r_i) \tag{35}$$

#### 3.2 Relative ratio

The expected relative ratio between the WT and the acapsular is:

$$R = \frac{q_w \exp(net_w t_w)}{q_a \exp(net_a t_w)\exp(net'_a(t_{tot} - t_w))} = \frac{K}{n_{0w} q_a \exp(net_a t_w)\exp(net'_a(t_{tot} - t_w))} \tag{36}$$

$$R = \frac{K}{n_{0w} q_a} \exp(-net_a t_w - net'_a(t_{tot} - t_w)) \tag{37}$$

Using the expression for $net'_a$ equation (32),

$$R = \frac{K}{n_{0w} q_a} \exp(-rrnet_w t_w - (net_a - rrnet_w) t_{tot}) \tag{38}$$

With $n_{0w}$ the initial number of WT bacteria, $n_{0w} q_w \exp(t_w net_w) = K$, leading to:

$$R = \frac{q_w^{rr}}{q_a} \left(\frac{K}{n_{0w}}\right)^{1-rr} \exp((rrnet_w - net_a) t_{tot}) \tag{39}$$

For the Oligo and LCM microbiota, the colonization process showed that the colonization probability was very similar for the WT and the acapsular, so that it can be assumed that $q_a = q_w$ in these cases. There is no reason *a priori* that it be different for GF mice, so this assumption also extends to GF mice. For SPF mice, the apparent colonization probability was quite different, so let us define $qq = q_a/q_w$. For the colonization process, as the *B.theta* population grows to larges sizes, $q_{i,1,app}$ the apparent colonization probability of type $i$ (WT of acapsular) in the colonization process (1 for 1 strain type), $q_{i,1,app} \simeq q_i(1 - c_i/r_i)$. Then:

$$qq_{SPF} = \frac{q_{a,1,app} r_w net_a}{q_{w,1,app} r_a net_w} \tag{40}$$

**When** $rr = ra/r_w = 1$:

Given that the WT and the acapsular grow at the same rate *in vitro*, the null assumption is to take then also growing at the same rate *in vivo*, but with different clearance rates, leading to different net growth rates. With this assumption,

$$R(rr = 1) = \frac{q_w}{q_a} \exp((net_w - net_a) t_{tot}) = \frac{1}{qq} \exp((net_w - net_a) t_{tot}) \tag{41}$$

$qq = 1$ for SPF, Oligo and LCM, and is $= \frac{q_{a,1,app} net_a}{q_{w,1,app} net_w}$ for SPF.

This prediction mixes results from the growth experiments ($net_w$ and $net_a$), and results from the colonization experiment ($qq = 1$ in most cases, $q_{a,1,app}$ and $q_{w,1,app}$ to calculate $qq$ in the SPF case).

**When $rr \neq 1$**

**Predicted relative ratio**

Reformulating (39):

$$R = \frac{q_w^{rr-1}}{qq}\left(\frac{K}{n_{0w}}\right)^{1-rr}\exp((rrnet_w - net_a)t_{tot}). \tag{42}$$

For GF, Oligo and LCM mice, $qq$ is taken as = 1, and for SPF, expression (40) is used; for $q_w = q_{w,1,app}/(1-c_w/r_w) = q_{w,1,app}(net_w + c_w)/net_w$. In summary, $qq$ and $q_w$ are taken from the colonization experiments, $net_w$ and $net_a$ from the growth experiments, $n_{0w}$ and $t_{tot}$ are controlled experimental parameters, $K$ is experimentally measured. As explained earlier, there are indications that $rr$ should be = 1, but in the confidence interval calculations we explore variations around 1.

**Relative ratio given the loss**

Reformulating (39):

$$R = \frac{q_a^{rr-1}}{qq^{rr}}\left(\frac{K}{n_{0w}}\right)^{1-rr}\exp((rrnet_w - net_a)t_{tot}) \tag{43}$$

As before, $qq$ is taken as 1, except for SPF mice, for which expression (40) is used.The difference with the previous expression is that now $q_a$ is taken from the competition experiment. There are 2 expressions linking $q_a$ to the apparent survival probability $q_{a,app}$.

- In the limit, when the population of acapsular remains high enough at the end of the experiment despite competition with the WT, then the same approximation as in the colonization experiments can be made, and $q_a \simeq q_{a,app}/(1-c_a/r_a) = q_{a,app}rr(net_w + c_w)/net_a$. Note that here one additional assumption is made, on the value of $c_w$. It is generally taken as the cecum turnover rate, which is its minimal value, but higher values are explored for the confidence intervals.
- The full expression is $q_a = q_{a,app}/(1 - g_{up}(g_{plat}(0)))$

#### 3.3 Predicted survival probability

The expression for the survival probability is $q_a(1 - g_{up}(g_{plat}(0)))$

The estimate for $q_a$ will be taken from the colonization experiment, with the approximation $q_{a,1,app} = q_a(1 - c_a/r_a)$, thus:

$$p_{surv} = q_{a,1,app}\frac{(1 - g_{up}(g_{plat}(0)))}{1 - c_a/r_a} \tag{44}$$

As seen earlier, $g_{up}$ depends on $net_a$, $t_w$, $r_a$, $c_a$. Also, $g_{plat}$ depends on $net'_a$, $t_{tot}$, $t_w$, $r_a c_w/r_w = rrc_w$, $c_a$. We note that $net'_a = net_a - rrnet_w$, that $r_a = rrr_w = rr(net_w + c_w)$ and that $c_a = r_a - net_a = rr(net_w + c_w) - net_a$. As $n_{0w}q_w\exp(t_w net_w) = K$, and $q_w \simeq q_{w,1,app}/(1 - c_w/r_w) = q_{w,1,app}(net_w + c_w)/net_w$, $t_w = \log(Knet_w/(n_{0w}q_{w,1,app}(net_w + c_w)))/net_w$.

So the survival probability ends up being dependent on $q_{a,1,app}$, $q_{w,1,app}$, (both measured in the colonization experiment), $net_a$, $net_w$ (both measured in the growth experiment), $t_{tot}$, $n_{0w}$ (both known controlled parameters of the experiment), $K$ (measured), $c_w$ (in general taken as its lower bound, the cecum turn over) and $rr$ (generally taken as 1).

#### 3.4 Effect of fixed delay in growth

If there was a delay $\tau_i$ for strain $i$ before starting growth after ingestion, then:

$$q_{i,1,app} = q_i\exp(-c_i\tau_i)(1 - c_i/r_i) \tag{45}$$

$K = n_{0,w}q_w\exp(-c_w\tau_w)\exp(net_w(t_w - \tau_w))$ (note that here $t_w$ is the time at which carrying capacity is reached, with duration $\tau_w$ of population decrease, and $t_w - \tau_w$ of net growth $net_w$ of the wild type), thus

$$\exp(net_w(t_w - \tau_w)) = \frac{K\exp(c_w\tau_w)}{n_{0,w}q_w} = \frac{Knet_w}{n_{0,w}q_{w,1,app}(net_w + c_w)} \tag{46}$$

The expected relative ratio between the WT and the acapsular is:

$$R = \frac{q_w \exp(-c_w \tau_w) \exp(net_w(t_w - \tau_w))}{q_a \exp(-c_a \tau_a) \exp(net_a(t_w - \tau_a)) \exp(net'_a(t_{tot} - t_w))} \tag{47}$$

As $net'_a = net_a - rrnet_w$, replacing $\exp(net_w(t_w - \tau_w))$ by previous expression, and after some calculations,

$$R = \frac{1}{qq} \left( \frac{K \exp(c_w \tau_w)}{n_{0w} q_w} \exp(net_w \tau_w) \right)^{1-rr} \exp((net_w + c_w)(rr\tau_a - \tau_w)) \exp((rrnet_w - net_a)t_{tot}) \tag{48}$$

If $rr = 1$, then:

$$R = \frac{1}{qq} \exp((net_w + c_w)(\tau_a - \tau_w)) \exp((net_w - net_a)t_{tot}) \tag{49}$$

As before, the assumption is $qq = 1$, except for the SPF, when:
$qq = q_{a,1,app}(1 - c_w/r_w) \exp(-c_w \tau_w)/(q_{w,1,app}(1 - c_a/r_a) \exp(-c_a \tau_a))$.

If $rr$ different from 1, in the case of a prediction from the rest of the experiments,

$$R = \frac{1}{qq} \left( \frac{K \exp(c_w \tau_w)}{n_{0w} q_w} \exp(net_w \tau_w) \right)^{1-rr} \exp((net_w + c_w)(rr\tau_a - \tau_w)) \exp((rrnet_w - net_a)t_{tot}) \tag{50}$$

$$R = \frac{1}{qq} \left( \frac{Knet_w}{n_{0w} q_{w,1,app}(net_w + c_w)} \exp(net_w \tau_w) \right)^{1-rr} \exp((net_w + c_w)(rr\tau_a - \tau_w)) \exp((rrnet_w - net_a)t_{tot}) \tag{51}$$

If $rr$ is different from 1, in the case of checking for internal consistency of the competition experiment,

$$R = \frac{1}{qq^{rr}} \left( \frac{K \exp(c_w \tau_w)}{n_{0w} q_a} \exp(net_w \tau_w) \right)^{1-rr} \exp((net_w + c_w)(rr\tau_a - \tau_w)) \exp((rrnet_w - net_a)t_{tot}) \tag{52}$$

with $q_a$ is such that $q_{a,app} = q_a \exp(-c_a \tau_a)(1 - g_{up}(g_{plat}(0))))$ (note that $g_{up}$ is for $t_{up} = t_w - \tau_w =$, and $g_{plat}$ for $t_{plat} = t_{tot} - t_w$).

And the expected survival probability is $q_a \exp(-c_a \tau_a)(1 - g_{up}(g_{plat}(0)))$, with $q_a \simeq q_{a,1,app} \exp(c_a \tau_a)/(1 - c_a/r_a)$, and thus

$$surv = q_{a,1,app}(1 - g_{up}(g_{plat}(0)))/(1 - c_a/r_a) \tag{53}$$

#### 3.5 List of parameters and values

| Symbol | Meaning (More info) | How determined / values taken (GF, Oligo, LCM, SPF) |
|---|---|---|
| $qq$ | $q_a/q_w$ (1) | 1, 1, 1, Depends on model |
| $n_{0w}$ | Number of WT bacteria in the inoculum (2) | Controlled and measured<br>$10^{7.63\pm0.1}$, $10^{7.54\pm0.17}$, $10^{7.31\pm0.49}$, $10^{7.56\pm0.20}$ |
| $K$ | Effective carrying capacity (3) | Measured<br>$10^{10.86\pm0.23}$, $10^{10.17\pm0.47}$, $10^{9.49\pm0.33}$, $10^{7.35\pm0.53}$ |
| $rr$ | Ratio of the growth rates, $r_a/r_w$ (4) | 1 [0.5-1.2] (for all microbiota) |
| $net_w$ | net growth rate, $r_w - c_w$ (5) | Measured in the growth experiments in Oligo and SPF<br>$0.53/h(0.39 - 0.69)$, $0.53/h(0.49 - 0.59)$, $0.53/h(0.49 - 0.59)$, $0.38/h(0.27 - 0.47)$ |
| $net_a$ | net growth rate, $r_a - c_a$ (5) | Measured in the growth experiments in Oligo<br>$0.40/h(0.29 - 0.52)$, $0.40/h(0.39 - 0.42)$, $0.40/h(0.39 - 0.42)$, $0.25/h(0.14 - 0.37)$ |
| $q_{a,app}$ | apparent survival probability (6) | Measured in the competition experiments<br>$10^{-1.27[-1.35,-1.19]}$, $10^{-3.4[-3.47,-3.33]}$, $10^{-2.63[-2.75,-2.51]}$, $10^{-6.59[-6.64,-6.54]}$ |
| $q_{a,1,app}$ | apparent survival probability (7)<br>when only acapsular | Measured in the colonization experiments (except GF)<br>$10^{-0.7[-1.3,-0.1]}$, $10^{-1.48[-1.65,-1.31]}$, $10^{-1.42[-1.58,-1.26]}$, $10^{-3.60[-3.77,-3.43]}$ |
| $q_{w,1,app}$ | apparent survival probability (7)<br>when only WT | Measured in the colonization experiments (except GF)<br>$10^{-0.7[-1.3,-0.1]}$, $10^{-1.54[-1.71,-1.37]}$, $10^{-1.49[-1.64,-1.34]}$, $10^{-2.34[-2.51,-2.17]}$ |
| $t_{tot}$ | Total experimental time (8) | fixed at 40h (taken 40h [40,48]) |
| $c_w$ | Loss rate of the WT (9) | Minimum is the cecum turnover rate<br>$0.13/h[0.13 - 0.23]$, $0.13/h[0.13 - 0.23]$, $0.13/h[0.13 - 0.23]$, $0.23/h[0.23 - 0.33]$ |
| $m_w$ | Mean growth delay for the WT (10)<br>Delay model only | 0h [0-4] |
| $m_a$ | Mean growth delay for the acapsular (10)<br>Delay model only | 6h [4-8] |
| $r_w$ | Growth rate of the WT | $r_w = net_w + c_w$ |
| $r_a$ | Growth rate of the acapsular | $r_a = rr(net_w + c_w)$ |
| $c_a$ | Loss rate of the acapsular | $c_a = r_a - net_a = rr(net_w + c_w) - net_a$ |
| $q_i$ | Survival probability initial bottleneck $i$ ($w$ or $a$) (11) | Estimated using $q_{i,1,app}$ or $q_{a,app}$ |
| $t_w$ | Time for the WT to reach carrying capacity | Estimated from the other parameters |

Detailed notes:

1. As the colonization experiment give very similar apparent probability for the WT and acapsular, except for the SPF, the assumption is that $qq = 1$ for all except SPF. For SPF, $qq = q_a/q_w$, and in the simple model, taking the approximation that in the colonization experiments, $q_{app} = q(1 - c/r)$, then $qq = (q_{a,app}(1 - c_w/r_w))/(q_{w,app}(1 - c_a/r_a))$, which can also be written as: $= (q_{a,1,app} net_w rr)/(q_{w,1,app} net_a)$.
2. Measured in the inoculum, averaged over the inoculum used for the given microbiota, ± the standard deviation between values for the different used inoculum (average and sd calculated on the log values). For GF, only one inoculum is used, so there is no standard deviation, the error is estimated ±0.1 of the log10 of the concentration.
3. Actually, what is measured is number of bacteria per g of cecum content, whereas $K$ is the absolute value. The assumption is that the cecum is about 1 gram. In reality, it is often a bit smaller, but the difference is small compared to the differences in observed final bacterial concentrations.
4. The rationale is that in vitro, they have the same growth rate, so there is no specific reason to believe they are different. So this ratio is usually taken as 1, and the confidence interval is taken as the minimal and maximal values from the fit from the growth rate.
5. We estimated the growth rate for acapsular strain in Oligo $net_a = 0.40/h(0.39 - 0.42)$ (the lower and upper bound are the lowest and highest fit values). For the WT strain in Oligo $net_w = 0.53/h(0.49 - 0.58)$, and in SPF $0.38/h(0.27 - 0.47)$. In other preliminary experiments, the growth rate of bacteria in Oligo and LCM was similar, so the same values are taken for Oligo and LCM. GF mice seem closer in terms of turnover to the LCM and Oligo mice, so the same growth rate is taken in GF for the WT strain. For the acapsular strain, we assumed lower clearance than in a colonized mice so we increased the $net_a = 0.46/h(0.39, 0.52)$. Also, we increased the window of uncertainty by $\pm 0.1/h$ to take into account the data limitation. For SPF, it is observed for the WT that the net growth rate is decreased by $0.15/h$, which is consistent with a higher turnover in the cecum of SPF mice (higher by 0.1/h compared to GF mice), which points towards an increase in $c$ rather than a decrease in $r$. For $net_a$ in SPF, in the absence of direct data, the assumption is to take the same decrease relative to the Oligo mice, with a higher uncertainty range to take into account the absence of direct data.
6. Measured from the tag loss in the competition experiment
7. Measured in the colonization experiments, the value taken is the estimate via the tag loss. For GF, a lower bound is the apparent survival probability in the competition experiment ($10^{-1.27} \simeq 10^{-1.3}$), as it is smaller than in the colonization experiments. An upper bound is considering that in the simple model, subsequent survival is at most equal to $(1 - c/r) = net/(net + c)$, and with a net growth rate of the order of $0.53/h$ (WT) and $0.40/h$ (acapsular) (both measure in Oligo but expected to be similar in GF), and the $c$ at least equal to $0.13/h$, the survival is at most $\simeq 10^{-0.1}$. The main value taken is intermediate (in log scale) between these 2 boundaries.
8. The initial survival probability may represent very early processes, in the stomach and small intestine, so the time spent in the cecum may be smaller than 48h, so it is why in exploring the parameter space the choice is made to take 40h-48h as the confidence interval.
9. The loss rate is at least equal to the cecum turnover rate. It is $0.13/h$ for the GF, and $0.23/h$ for the SPF. LCM and Oligo are thought to be closer to the GF. If would be higher in the presence of killing. Without evidence of killing, the main value is this minimal value. The upper bound is taken as this value, $+0.1/h$ to represent the impact of potential killing
10. The delay model explores the possibility that the acapsular takes a longer time to recover and grow back. In the growth curves, all are consistent with exponential growth from 12h onwards, with points beforehand that suggest some delay, though exact quantification is difficult. In terms of results, what still matters is the difference between the WT and the acapsular. We choose 0h and 4h as the typical delay for the WT vs. the acapsular, with boundaries [0-5h], [0-14h] respectively. In a second model of delay, we explore the effect of delay spread. For the WT, the main value chosen for the standard deviation is very small (0.01h), to represent the situation of no delay (but not zero, to avoid numerical issues), with [0.01-3] as the boundaries. For the acapsular, we assume a larger spread, of typically 2h, interval [1h-8h].

11. In the colonization experiments, the bacterial lineages reach and remain at a large population, so later loss is negligible, thus $q_{i,app} \simeq q_i(1 - c_i/r_i)$ for the simpler model. See text for other models.

##### 3.6 Computation

The R code is attached in the supplementary material. The results are first computed for the main values of the parameters, for all the models. Then, a chosen number of iterations (in general 1000), a new set of parameters is taken at random. For each parameter, with probability 0.5 it is chosen randomly uniformly between the lower bound and the main value; and with probability 0.5 it is chosen uniformly between the main value and the upper bound of the confidence interval. It is checked that these parameters lead to $q_a$ and $q_w$ as calculated from the expression from all models and $q_{i,1,app}$ is below 1. For the simple model, if it does not work, all the parameters are chosen again at random. For the other model, first, for at most 300 iterations, only the parameters linked to the delay are changed, and then if the result gives still survival greater than 1, a full set of parameters is again chosen randomly. This procedure help to avoid bias in the choice of the parameters value.

The results are computed for each of these sets of parameters, and the confidence interval is given as the mean $\pm$ the standard deviation on all these iterations.

### 4 Acute challenges

#### 4.1 Estimate of $n_0$

We need to estimate the number of tagged bacteria in the cecum at the start of the challenge from the intermediate data collected in feces, which are a subsampling of the cecum, with potential bias.

##### 4.1.1 Cecum mass

A first question is the mass of the cecum, as the feces gives only access to a concentration. For all cases, the conditions are like the control conditions up to day 0, thus reasonably at day 0 the cecum is expected to weight like the cecum in the control case at day 3 at the end of the experiment. The average amount of sampled material from the cecum is 0.66 grams, standard deviation 0.11g.

##### 4.1.2 Concentration in feces vs. cecum

At day 3, the data consists of both feces and cecum, thus concentrations can be compared, see Figure 6. Using only the data from the untagged strain (less bias because no missing points, but overall there is no large difference between dashed and dotted lines), feces are $10^{0.67+-0.23}$ more concentrated than the cecum for the control ($\pm$ represent twice the standard deviation for a 95% confidence interval), feces are slightly less concentrated than the cecum for high-fat diet (HFD), and feces are less concentrated than the cecum in attenuated Salmonella infection (Stm).

##### 4.1.3 Factor to convert concentration in feces at day 0 to absolute number in cecum at day 0

To estimate $n_0$: The total number of tagged strain in a mouse at day 0 is taken as the number of erythromycin-resistant colonies in feces, divided by the weight of the sample to obtain the concentration, then multiplied by the mean cecum mass. Finally, the concentration factor for the control case is applied. Then for a given tag, the counts are used to get the proportion of that tags in the total number of tags. We estimate errors by applying this method to the data in feces at day=3 and comparing with the cecum at day=3. Our estimate gives the order of magnitude, but interval of confidence is divided by 4 or multiplied by 4, to account for the variation in the relation between feces and cecum concentrations.

##### 4.1.4 Tags unseen at day 0

In most cases, tags are both seen in feces at day 0, and in cecum at day 3.

In a fraction of cases, tags are seen in feces at day 0, but not in cecum at day 3. In this case, as the sample is a significant portion of the cecum, it is unlikely that the tag is actually still present, so here we count this situation as a loss.

In the counts of tagged bacteria, there are a few cases where some tags are not seen at day 0, but still are seen at day 3, due to the feces sample weighting in average more than 10 fold less than the cecum and these tagged bacteria being in very low numbers.

This also points to the possibility that in the cases where the tags are not seen neither at day 0 nor at day 3, they may have been actually present at day 0 and lost before day 3.

For the loss, we will analyze the data:

- Excluding all cases in which the tags were not seen at day 0.
- Excluding all cases in which the tags were not seen neither at day 0 nor day 3, and for tags not seen at day 0 and seen at day 3, $n_0$ for day 0 is taken as the average estimated total number of tags in that mouse at day 0, divided by 6 (total number of tags)
- For all cases with no tags seen at day 0, $n_0$ for day 0 is taken as the average estimated total number of tags in that mouse at day 0, divided by 6 (total number of tags)

In practice, the differences in the results are negligible.

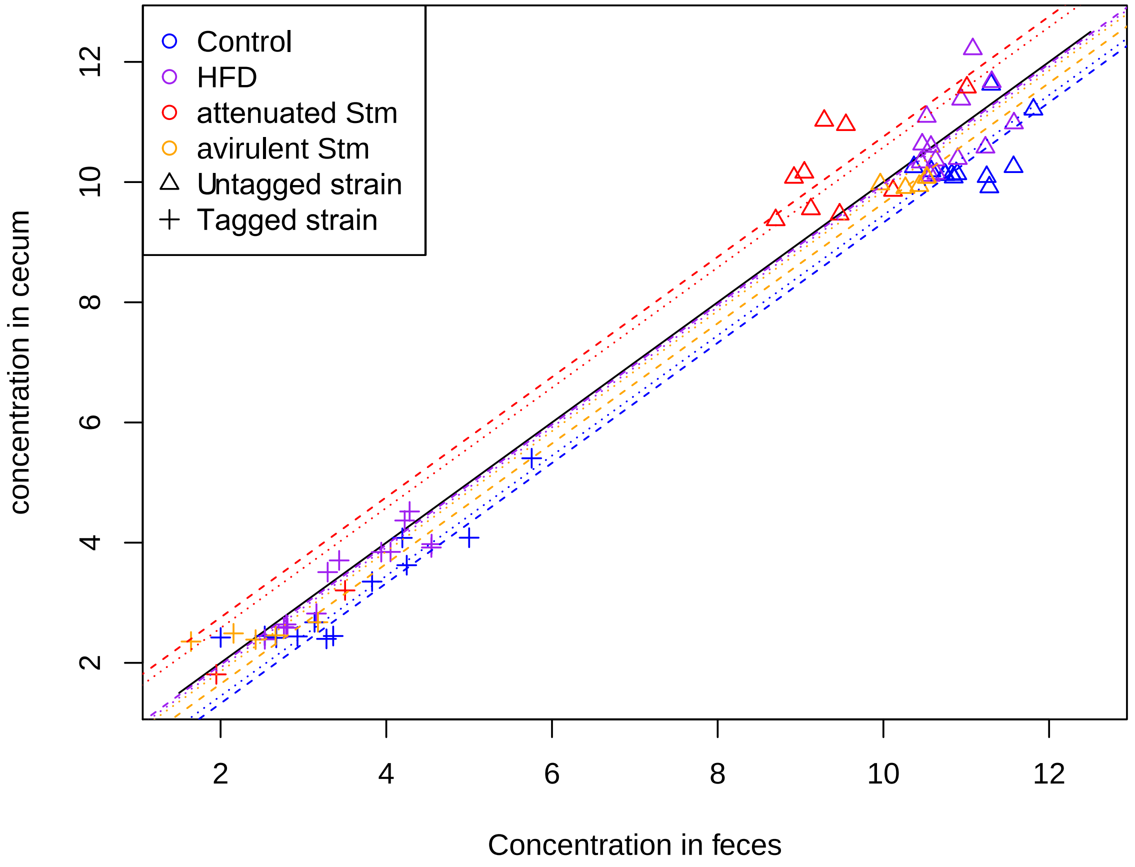

Figure 6: Concentration in bacteroides (all (Tet), and tagged (Ery)) in the cecum relative to the feces, at day 3, in the challenge experiments. The dashed lines represent the average ratio taking into account only the Tet data, whereas the dotted lines represent the average taking into account all the data, except the case of total loss of Ery (always with salmonella), which may bias the data.

#### 4.2 Method of the loss

##### 4.2.1 Survival probability of a bacterial lineage

We use the same method as in previous section to estimate the colonization probability, except that $n_0$ is estimated from the feces at day 0 instead of the inoculum.

There was no loss for neither the control nor HFD (there is one loss in HFD, but it is also a loss at day=0, for a large total number of tagged strains in the feces at day 0 in that mouse, so it is likely that the tag was actually lost at day 0). Then to obtain a bound, the case with the smallest $n_0$ is taken as lost and the survival probability is calculated from it.

##### 4.2.2 Results

- There is no loss for the control case, so the best estimate of $\beta$ is 1, with lower bound $10^{-1.5}$.
- There is no loss for the HFD case, so the best estimate of $\beta$ is 1, with lower bound $10^{-1.75}$.
- For attenuated Stm, $10^{-3.28\pm0.10}$ (for the incertitude: for the log10 of the concentration, the standard error is 0.21, but there are 14 different mice for Stm, thus 0.06 overall expected on the combined data if the ratio of the concentrations are independent in the different mice; for the log10 of the survival probability using a Bayesian approach, the standard deviation is found to

be around 0.08. So overall the standard deviation is expected to be around 0.10, then taking 2 standard deviations we find the result)

- For the avirulent Stm, $10^{-2.35 \pm 0.3}$ (similar reasoning)

Thus attenuated Stm imposes a relatively strong bottleneck on *B.theta*. The bottleneck from the avirulent Stm is less stringent. HFD does not have a strong enough effect to be detected in the experimental conditions.

##### 4.2.3 Interpretation

This survival probability gives the probability that a bacteria present at day=0 has still a lineage in the cecum at day 3.

It could either come from a temporary bottleneck, or from constant loss during the challenge. If there is a constant turnover rate $c$, then the survival after a time $t$ is $1/(1+ct)$. For $c$ in GF mice of 0.13/h, and for 3×24h, we find a survival of about 0.1 ($10^{-1}$), which is compatible with the results for the Control and HFD.

HFD could be causing a larger $c$, but as no loss is observed, we cannot quantify it.

For Stm, the decrease in total population size is at most a factor of 10, whereas survival probability is well below 0.1, pointing to a more stringent bottleneck followed by re-growth (and possibly additional continuous loss).
